# Characterization and molecular insights of a chromium-reducing bacterium *Bacillus tropicus*

**DOI:** 10.1101/2024.09.19.613096

**Authors:** Shanjana Rahman Tuli, Md. Firoz Ali, Tabassum Binte Jamal, Md. Abu Sayem Khan, Nigar Fatima, Irfan Ahmed, Masuma Khatun, Shamima Akhtar Sharmin

**Affiliations:** Environmental Biotechnology Division, National Institute of Biotechnology, Ganakbari, Ashulia, Savar, Dhaka 1349, Bangladesh; Department of Biotechnology and Genetic Engineering, Mawlana Bhashani Science and Technology University; Department of Microbiology, University of Dhaka

**Keywords:** *Bacillus tropicus*, chromium-reducing bacteria, bioremediation, heavy-metal tolerant, whole genome sequencing

## Abstract

Environmental pollution from metal toxicity is a widespread concern. Certain bacteria hold promise for bioremediation that converts toxic chromium into a less harmful form, promoting environmental cleanup. In this study, we report the isolation and detailed characterization of a highly chromium-tolerant bacterium, *Bacillus tropicus* CRB14. The isolate is capable of growing on 5000 mg/l Cr (VI) in LB agar plate while on 900 mg/l Cr (VI) in LB broth with an 86.57% reduction ability within 96 hours of culture. It can also tolerate high levels of As, Cd, Co, Fe, Zn, and Pb. The plant growth-promoting potential of the isolate was demonstrated by a significant activity of nitrogen fixation, phosphate solubilization, IAA, and siderophore production. Whole-genome sequencing revealed that the isolate lacks plasmids for Cr resistance, suggesting genes reside on its chromosome. The presence of the *chrA* gene points towards Cr (VI) transport, while the absence of *ycnD* suggests alternative reduction pathways. The genome harbors features like genomic islands and CRISPR-Cas systems, potentially aiding adaptation and defense. Analysis suggests a robust metabolism, potentially involved in Cr detoxification. Notably, genes for siderophore and NRP-metallophore production were identified. Whole Genome Sequencing (WGS) data also provides the basis for molecular validation of various genes. Findings from this study highlight the potential application of *Bacillus tropicus* CRB14 for bioremediation while plant growth promotion can be utilized as an added benefit.

## Introduction

Chromium (Cr) is one of the common pollutants released from various industries, including steel production, electroplating, tanning leather, wood processing, the production of Cr pigments, the manufacture of glass, ceramics, and cement, etc. Among these, leather processing industries are one of the highest contributors to Cr pollutants in Bangladesh [1]. In order to turn animal hides into leather, almost all leather tanneries employ chemicals that include metals. The wastewater that these businesses release into the environment is one of the main known sources of metal pollution [2]. Approximately 55-70% of the chromium salt that enters the tanning fluid during tanning is fixed in the leather, with the remaining portion ending up in the effluent. The tanning float produces the majority of the chrome effluent, with a sizable part coming from the sammying and re-tanning operations as well. While chromium concentrations in spent liquor range from 2000 to 5000 ppm [3]. International bodies within the EU, such as the Helsinki Commission and the Oslo-Paris Convention, have issued recommendations on chromium discharge levels. The maximum discharge limit to the aquatic environment in the EU is 1 and 5 mg L^−1^ for Cr^VI^ and Cr^total^, respectively [4]. It is conceivable to bring the chromium concentrations in the tannery waste stream down to acceptable levels, but doing so would require significant investment and burden the tanneries with additional operating costs [5].

The toxicity of Cr is significantly dependent on its chemical form. Hexavalent Cr compounds are extremely toxic, mutagenic, and carcinogenic [6]. This is because Cr (VI) is more quickly and actively transported through biological membranes as a result of its high solubility, bioavailability, and oxidizing qualities [7]. It is also widely known that some species require a tiny quantity of Cr (III) for both growth and metabolic potential [8], [9]. Cr (III) in the environment becomes insoluble at neutral or alkaline pH levels and forms a precipitate that is easily removed from aqueous media. However, trivalent Cr (Cr III) is 100 times less hazardous than Cr (VI). Asthma, a perforated nasal septum, renal failure, dermatitis, hepatitis, and other conditions can make human organs like the skin, liver, kidneys, and respiratory tracts more susceptible to Cr (VI) toxicity. Cr (VI) is more stable in alkaline pH ranges [10], [11]. Consequently, the strategy of converting Cr (VI) to Cr (III) in industrial wastes is considered very useful in reducing immediate toxicity.

The toxicity of Cr is significantly dependent on its chemical form. Hexavalent Cr compounds are extremely toxic, mutagenic, and carcinogenic [6]. This is because Cr (VI) is more quickly and actively transported through biological membranes as a result of its high solubility, bioavailability, and oxidizing qualities [7]. Asthma, a perforated nasal septum, renal failure, dermatitis, hepatitis, and other conditions can make human organs like the skin, liver, kidneys, and respiratory tracts more susceptible to Cr (VI) toxicity and it is more stable in alkaline pH ranges [10], [11]. While Cr (III) in the environment becomes insoluble at neutral or alkaline pH levels and forms a precipitate that is easily removed from aqueous media. It is also widely known that some species require a tiny quantity of Cr (III) for both growth and metabolic potential [8], [9]. However, Trivalent Cr (Cr III) is 100 times less hazardous than Cr (VI). Consequently, the strategy of converting Cr (VI) to Cr (III) in industrial wastes is considered very useful in reducing immediate toxicity.

To lessen the toxicity of Cr in the environment, tests are conducted on physical, chemical, and biological processes. These techniques usually prove costly and inefficient for widespread use. Numerous bacteria possess the ability to convert Cr (VI) into Cr (III), making this bioremediation method more viable, cost-effective, and long-lasting while achieving the highest level of remediation [12], [13], [14].

From chromium-contaminated tannery sites, several Cr-resistant bacteria, including *Bacillus* species, have been isolated [15], [16], [17], [18]. In Cr-resistant bacteria, both enzymatic and non-enzymatic reduction of Cr (VI) to Cr (III) has been shown [19]. Additionally, some of these bacteria exhibit beneficial characteristics toward plants, such as the promotion of plant growth (PGP). It was discovered that the combination of PGPB and particular contaminant-degrading bacteria performed well [20]. PGP has been studied for nitrogen fixation, siderophore production, phosphate solubilization, ammonia, and indole-3-acetic acid synthesis. Therefore, these bacteria have the potential to be used in sustainable agricultural production, particularly in contaminated areas [21]. The rate and efficiency of Cr (VI) reduction can be influenced by the initial concentration and environmental factors such as pH, temperature, and the presence of nutrients can affect the activity of chromium-reducing microbes. Different microbial strains have varying capacities to reduce Cr (VI). Nonetheless, a considerable knowledge vacuum persists regarding the precise mechanisms behind these actions. Therefore, this mechanism can be utilized for bioremediation and environmental cleanup if we have a better understanding of how bacteria achieve this Cr reduction and the factors impacting it.

Whole-genome sequencing (WGS) has become an effective approach that allows bacterial genome study and functional difference comparison with the current information [22], [23]. This anticipates the related pathways which helps in identifying the genes involved in Cr detoxification. This study set out to characterize in detail a strain of bacteria known as *Bacillus tropicus* CRB14, which was isolated from a local tannery effluent and is resistant to several heavy metals, including Cr. We investigated its resistance to Cr (VI), reduction potentials, tolerance to other heavy metals, growth-promoting qualities, growth-promoting effects of pH and temperature, and biofilm formation. To our knowledge, no study has been conducted to explore the Cr remediation mechanisms of *B. tropicus*. This study enhances the use of *B. tropicus* CRB14 in the detoxification of heavy-metal-contaminated locations by providing a rational basis for understanding chromium resistance and reduction through biochemical and genomic evidence.

## Materials and methods

### Collection and processing of Sediments

Tannery sediment was collected from the Hazaribagh tannery area of Dhaka city in sterile bottles and immediately transferred to the laboratory. Samples were diluted in normal saline and inoculated (0.1 ml) on Luria Bertani (LB) agar plate containing 100mg/l of potassium dichromate (K_2_Cr_2_O_7_) as Cr(VI) by spread plate method. Plates were incubated at 35℃ for 5 days. Useful twenty-five single colonies were selected and sub-cultured onto fresh LB agar medium and preserved at −80C in glycerol stock for further studies. After that, from the stock, we took randomly five isolates for further studies. **Figure 01** provides a schematic overview of the entire study.

**Figure 01:**
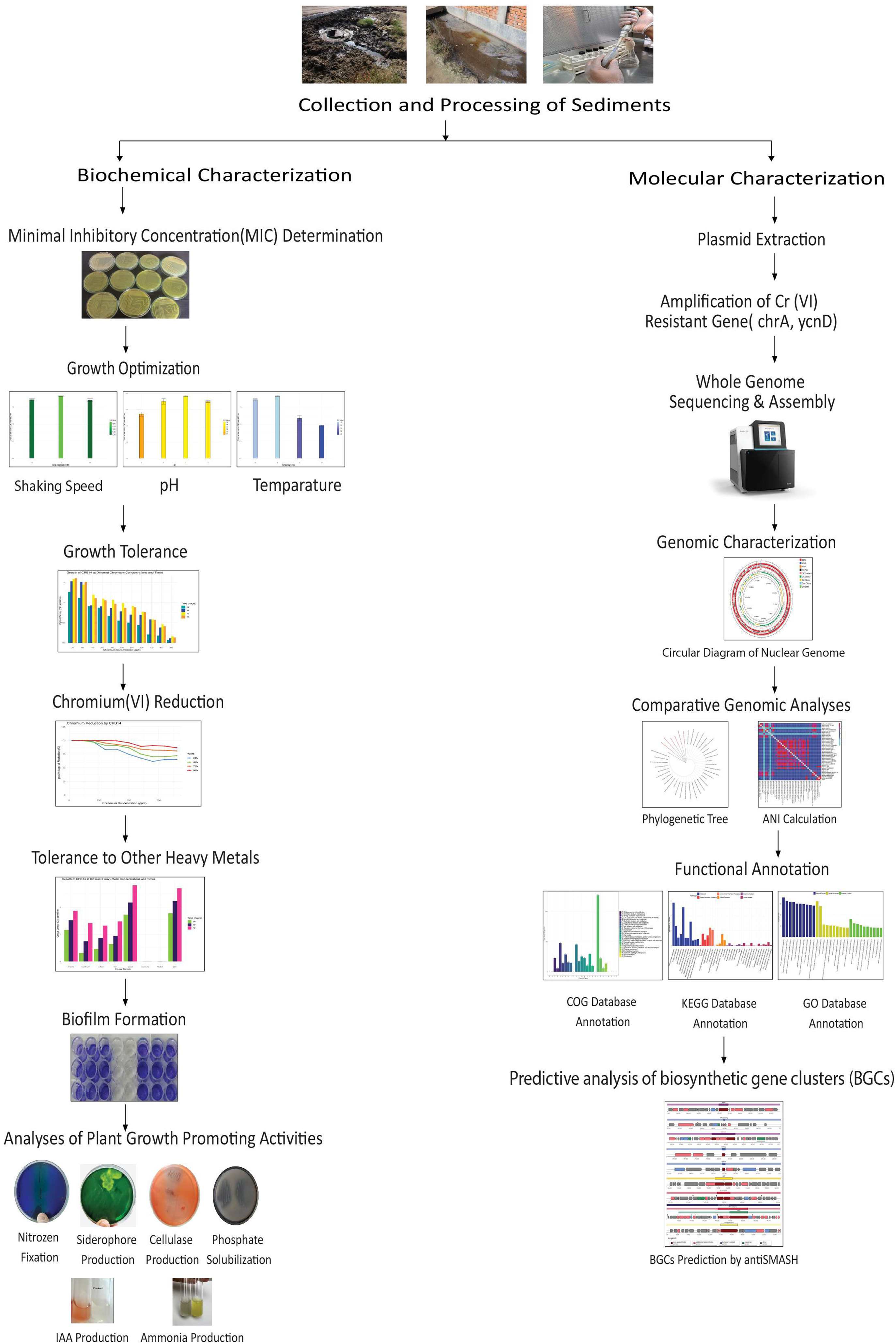
Graphical representation of the complete workflow for characterizing the chromium-reducing bacterium CRB14, starting with bacterial isolation, growth tolerance, and PGP analyses, and progressing through genome sequencing, functional annotation, and comparative analysis.

### Minimum inhibitory concentration (MIC) Determination

MIC of Cr (VI) for the isolates was determined on LB agar plates supplemented with filter-sterilized Cr (VI) at different concentrations (100 mg/L to 6000 mg/L) [24]. After incubation at 35 °C, for 5 days, the highest concentration of Cr (VI) which permitted growth and beyond which there was no growth was considered as MIC of Cr (VI) for the isolates tested.

### Optimization of physical factors (pH, temperature, shaking speed)

To identify the best growth condition, we incubated the bacteria in LB broth with 100 mg/l Cr (VI) at different pH (6,7,8,9), temperatures (30°C, 35°C, 37°C, 40°C) and shaking speeds (120rpm, 150rpm, 180rpm).

### Growth Tolerance to Chromium (VI)

Growth tolerance was performed by growing the isolate in 50 mL LB broth supplemented with 0, 25, 50, 100, 200, 300, 400, 500, 600, 700, 800, 900, 1000 mg/L of Cr (VI) as described by R Baldris et al. (2018) [19].

### Chromium (VI) Reduction Assay

To investigate the chromium (VI) reduction efficiency Cr(VI) reduction assay was performed by growing the isolate in 50 mL LB broth supplemented with 0, 25, 50, 100,200, 300, 400, 500, 600, 700, 800, 900,1000 mg/L of Cr (VI). A 0.5 mL aliquot of the original culture (OD 600 = 1.0±0.05) was transferred to each of a series of 50 mL aliquots of LB medium and the mixtures were incubated at 35℃, pH 8.0 with shaking at 150 rpm for 96hr to investigate the abilities of the isolates to reduce Cr (VI) as described by R Baldris et al. (2018) [19].

### Tolerance to other heavy metals

The effect of the various metals (Cd, Hg, Ni, Co, Fe, Zn, and Pb) on the growth of bacterial isolate was examined in LB broth (5 mL) supplemented with 100 mg/L of selected metals and 50 mg/L Cr (VI). The stock solutions were prepared from the analytical grade salts HgCl2, NiCl2-2H2O, ZnCl2, CuCl2, NaAsO2, PbSO4, CdCl2, FeSO4 for Hg (II), Ni (II), Zn (II) and Cu(II), As (III), Pb(II), Cd(II), Fe(II) ions. The tubes were inoculated with 25 μL (OD 600 = 1.0 ± 0.05) of the inoculum and incubated at 35 °C for 72hr. The growth of the bacterial isolate was measured spectrophotometrically at 600 nm [25].

### Morphological and biochemical characterization

The bacterial isolate CRB14 was identified by morphological and biochemical parameters by adopting the standard method [26]. Morphological characterization was done from pure culture of the isolate grown on LB medium agar at about 35°C under ambient conditions. The isolate was examined for its size, shape, margin, consistency, elevation, pigmentation, Gram reaction, and cell morphology. A presumptive identification was performed by the following tests: Gram staining, catalase test, oxidase test, urease test, indole test, methyl red-Voges Proskauer (MR-VP) test, citrate test, starch hydrolysis and nitrate reduction test, carbohydrate utilization test, motility test.

### Microtiter plate assay (quantitative assays for biofilm formation)

A crystal violet staining method was employed to examine the biofilm-forming abilities of the isolate with some modifications. Bacteria were first grown overnight in LB broth with or without 25, 50,100, and 200 mg/mL chromium VI as described by Baldris et al. (2018) [19].

### Analyses of plant growth-promoting activities

#### Production of indole acetic acid (IAA)

The amount of IAA produced by the isolate was determined by following the method described by Gordan and Weber (1951) [27]. Initially, the bacterial isolates were cultured in LB medium (∼108 CFU/ml) and culture was taken afterward to continue the assay. Using the standard curve for IAA, the amount of IAA produced by each inoculation was calculated.

#### Phosphate solubilization test

The inorganic phosphate solubilization ability of the bacteria was observed qualitatively by spot inoculation of the bacterial isolate on the NBRIP growth medium [28]. The formation of transparent halo zones around the bacterial colonies after 5 days of incubation at 35°C was considered as an indication of phosphate solubilizing activity. For the quantitative assay, a 100-μl aliquot of bacteria (∼108 CFU/ml) was quantitated by the molybdenum blue method in NBRIP broth [29]. The quantitative data about the amount of P-solubilized was extrapolated from the standard curve.

#### Nitrogen fixation assay

To determine the nitrogen-fixing capability of the isolate, 100 μl of bacterial inoculum (∼108 CFU/ml) was inoculated into a petri plate and conical flask containing 50 ml nitrogen-free New Fabian Broth (NFB) media and incubated for 24 hours at 35°C at 150 rpm on a shaker. The amount of fixed atmospheric nitrogen was determined by the Kjeldahl method [30].

#### Siderophore production test

Detection of siderophore production was done by following Schwyn and Neiland’s universal chrome azurol S (CAS) method [31]. Both qualitative and quantitative methods were used to estimate the siderophore production by the bacterial strain as described by Arora & Verma (2017) [32].

#### Cellulase Production Test

Pure cultures were screened for cellulase activity by plating on Congo Red agar media [33]. Bacterial isolate was streaked on Congo Red agar media and incubated at 35°C for 48 hours.

The use of Congo-Red as an indicator for cellulose degradation in an agar medium provides the basis for a rapid and sensitive screening test for cellulolytic bacteria. Colonies showing discoloration of Congo-Red were taken as positive cellulose-degrading bacterial colonies.

#### Ammonia Production

To detect ammonia production, bacterial isolate was grown in peptone broth and incubated at 37°C for 48 hours. After incubation, 0.5 ml of Nessler’s reagent was added to the bacterial suspension. The development of brown to yellow color indicated ammonia production [34].

### Molecular Characterization

#### Plasmid extraction

According to the manufacturer’s instructions, plasmid isolation was carried out using a commercial kit (Pure Plasmid Mini Kit, Cowin Biotech Co., Ltd, China). Plasmid was visualized with UV light following electrophoresis on 0.8% agarose gel stained with ethidium bromide (0.5 mg/mL).

#### DNA extraction and amplification of Cr (VI) resistant gene

Genomic DNA was extracted using Invitrogen TM Pure Link TM Genomic DNA Mini Kit (Catalog no. K182001; Life Technologies Corp., Carlsbad, CA 92008, USA) following the manufacturer’s protocol. The amplification of the chromate reductase gene was performed by PCR amplification with the primers listed in **Table 01**. PCR reactions were conducted in 25 microlitre aliquots containing 50 ng template DNA, 10 pM of each primer, GoTaq® Green Master Mix, and nuclease-free water. The reaction mixture was subjected to denaturation at 95°C for 5 min, followed by 35 cycles consisting of denaturation at 95°C for 30s, annealing at 54°C and 53°C (respectively) for 45s, and extension at 72°C for 2 min and a final extension of 72°C for 10 min. The PCR products were separated on 0.8% agarose gels in TBE buffer stained with ethidium bromide (0.5 mg/mL) and visualized with UV light.

**Table 01:**
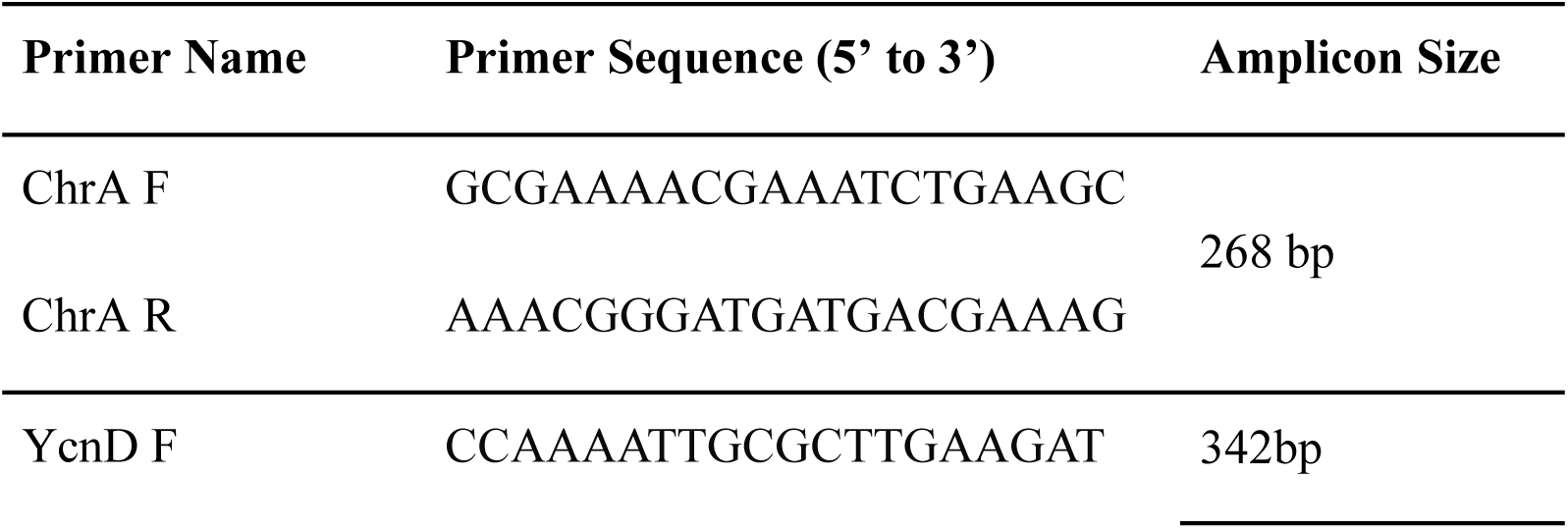
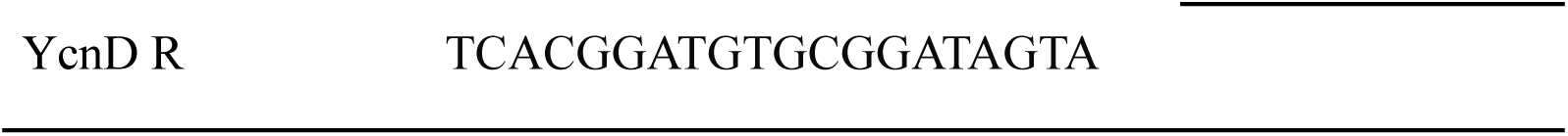
Primers for PCR amplification.

#### Whole genome sequencing and assembly

The DNA library was prepared using standard Illumina DNA prep workflow, which consists of tagmentation of genomic DNA, post-tagmentation cleanup, amplification of tagmented DNA, library cleanup, and library pooling steps. DNA prep insert size was 600 bp. Afterward, whole genome sequencing was performed in NovaSeq 6000 Illumina platform (2×150-bp paired-end reads) using NovaSeq 6000 Reagents (300 cycles) (Illumina, San Diego, CA, USA). The quality of raw sequencing data (.fastq files) was assessed using FastQC v 0.11.8, ensuring reads with high quality (Phred score > 30) [35]. Clean reads were obtained by filtering out adapters and low-quality reads from the processed data using Trimmomatic (version v0.39), followed by de novo assembly using SPAdes v4.0.0 [36], [37].

#### Genomic components

Prokka v1.12 and RAST tools were employed for predicting coding genes, tRNAs, and rRNAs [38]. A genomic circular map of the strain was generated using the Proksee web server, integrating GC ratio, GC-skew, and genome sequencing depth data [39]. Genomic islands (GIs) were predicted utilizing the IslandPath-DIOMB GI prediction method of the Island Viewer online tool (version 4.0) [40]. Clustered regularly interspaced short palindromic repeats (CRISPR) loci and antibiotic resistance gene (ARGs) were identified using the CARD Resistance Gene Identifier v1.2.1 tool, utilizing the Proksee web server [41], [42].

#### Genome identification and comparison

The 16S ribosomal DNA (rDNA) gene sequence was annotated within the genome, and its homology was assessed by comparing it with 16S rRNA gene sequences from other strains available in the GenBank database [43]. The multiple sequence alignment tool MAFFT was utilized for sequence alignment and a phylogenetic tree was constructed from the 16S rDNA using the maximum likelihood method [44]. The resulting tree was then visualized and annotated using the Interactive Tree Of Life (iTOL) version 6.9 [45]. Additionally, the average nucleotide identity (ANI) was calculated using the FASTANI tool on the Galaxy platform (Galaxy Version 1.3) [46].

#### Functional annotation

An in-depth annotation of predicted coding protein sequences was conducted by comparing them with various databases including Clusters of Orthologous Groups of proteins (COG), Kyoto Encyclopedia of Genes and Genomes (KEGG), Gene Ontology (GO), RefSeq, Pfam, and SwissProt [47], [48], [49], [50], [51], [52]. COG annotation was performed using eggNOG-mapper v2 to assign putative functions to proteins by identifying orthologous groups across different species based on evolutionary relationships [53], [54]. Furthermore, to match predicted genes with the KEGG database, BlastKOALA was employed, enabling the identification of genes involved in specific biological pathways via KEGG Orthology (KO) numbers obtained from blast alignment results [55], [56]. GO annotation was performed to evaluate the molecular functions of individual gene products, their active cellular components, and the processes they contribute to. GO was annotated using the Gene Ontology web server [48].

#### Prediction of biosynthetic gene clusters (BGCs)

The antibiotics and secondary metabolite analysis shell (antiSMASH) online version 7.1.0 was used to identify potential biosynthetic gene cluster (BGC) regions, and the resulting job ID was submitted to the biosynthetic gene cluster family (BiG-FAM) database to obtain the corresponding Gene Cluster Family (GCF) classification [57], [58]. The antiSMASH analysis was conducted using fasta DNA as input, with a relaxed detection strictness setting.

## Results

### Minimal Inhibitory Concentration (MIC)determination of the isolates

MIC of the bacterial isolate to Cr (VI) was assessed by the growth response of the strains under 100-6000 mg/l Cr (VI) concentrations for five days. Among the five tested isolates only one isolate CRB14 was able to grow up to 5000 mg/l Cr (VI). Therefore, we chose this isolate for further studies.

### Optimization of physical factors (pH, temperature, shaking speed)

After incubation of the isolates in LB broth with 100 mg/l Cr (VI) at different pH, temperatures, and shaking speeds we found out that the bacteria grew well at 35°C with 150 rpm shaking at pH 8 (Supplementary Figure 01).

### Growth tolerance to Chromium (VI)

Bacterial strain CRB14 was exposed to 25-1000 mg/l of Cr (VI) in LB broth for growth tolerance study. The isolate was found to grow up to 900 mg/l of Cr (VI), although growth reduction was observed with increasing concentration of chromium. The bacterial density (OD600) with 25-1000 mg/l Cr(VI) in LB broth was measured after 24hr, 48hr, 72hr, 96hr of culture. Bacterial strain CRB14 showed better growth on Cr(VI) concentration from 25-500 mg/l, a constant decrease in growth of CRB14 was observed at higher concentrations, and no growth was found above 900 mg/l of Cr(VI) (**Figure 02**).

**Figure 02:**
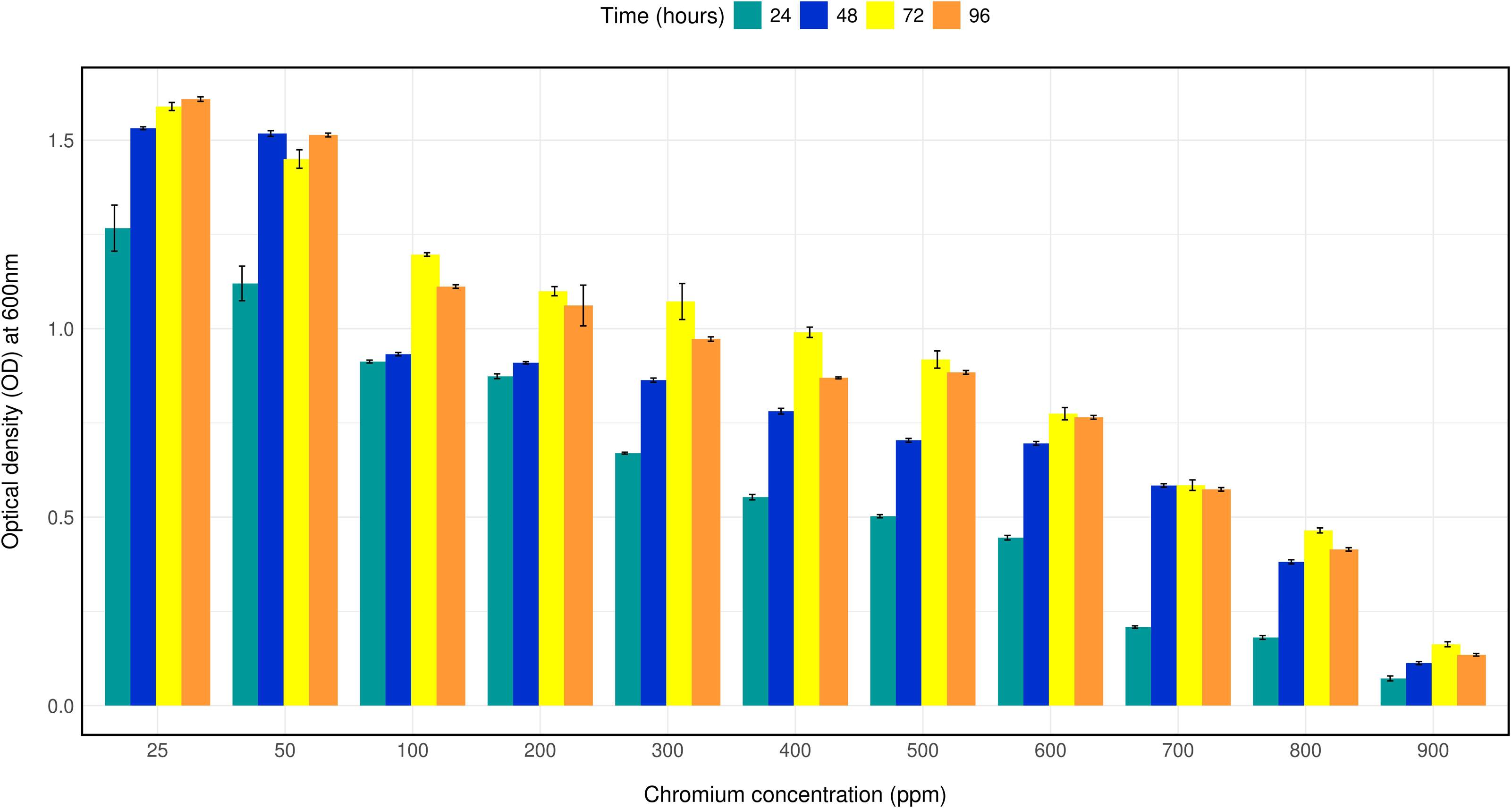
Bar plot depicting the growth of isolate CRB14 at different concentrations of Cr. The cells were cultured on Luria–Bertani broth supplemented with 25,50,100, and 200,300,400,500,600,700,800,900 mg/L Cr (VI), respectively. The optical density was measured after incubation for 24 h, 48h, 72hr, and 96h at 35°C.

### Chromium (VI) Reduction Assay

The bacterial strain CRB14’s capacity to reduce chromium (VI) was evaluated at concentrations ranging from 25 to 900 mg/l. As can be observed in Fig 2, at 900 mg/ml Cr (VI) concentrations, it is measured that 65.2%, 72.11%, 80.73%, and 86.57% Cr (VI) reduced after 24 hours, 48 hours, 72 hours, and 96 hours of culture, despite the isolate’s reduction effectiveness steadily decreasing with concentration (**Figure 03**).

**Figure 03:**
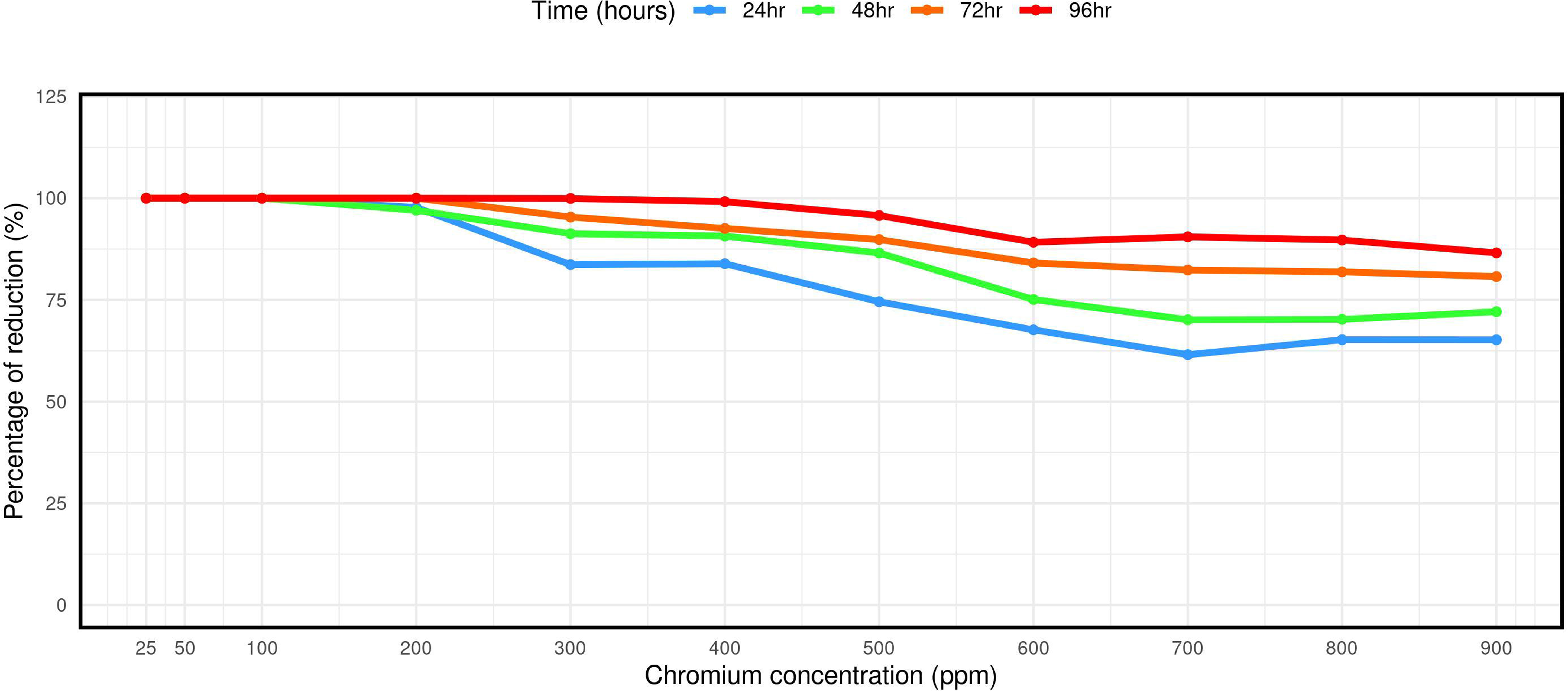
Reduction of Cr (VI) by isolate CRB14. The cells were cultured on Luria Bertani broth supplemented of 0, 25, 50, 100, 200, 300, 400, 500 600, 700, 800 and 900 mg/L Cr (VI). The Cr (VI) reduction activity was measured after incubation for 24 h, 48 h, 72 h, 96h and at 35°C.

### Tolerance to other metals

To observe the growth tolerance to other heavy metals the isolate was cultured in LB broth containing 100 mg/L of the salt of the selected metals along with 50 mg/L Cr (VI). The isolate was able to grow in cadmium, cobalt, iron, zinc, lead, and arsenic salts after twenty-four hours of culture. On the other hand, no growth was observed in the presence of nickel and mercury salts after seventy two hours of culture (**Figure 04**).

**Figure 04:**
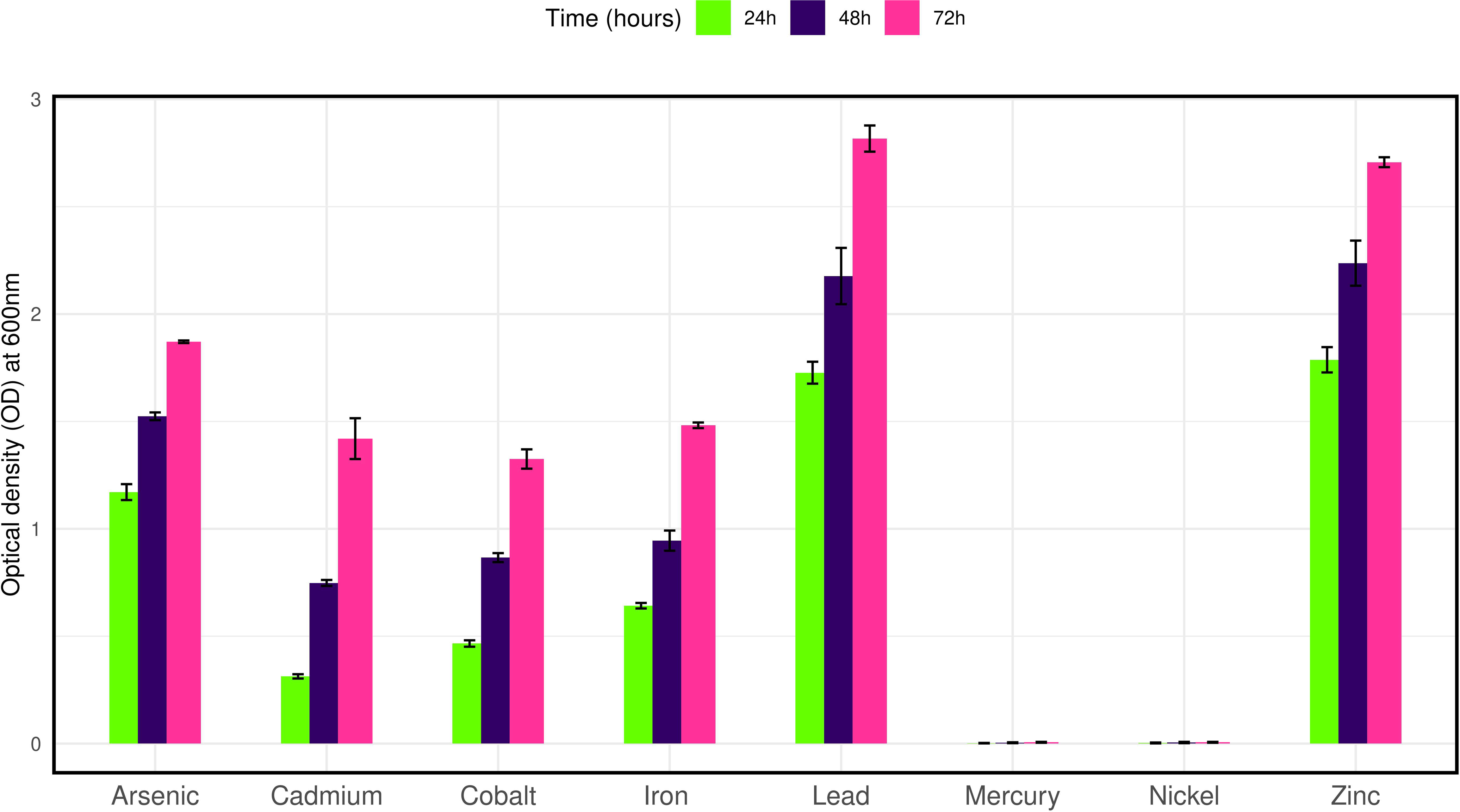
Bar plot representing the growth of isolate CRB14 in the presence of various heavy metals.

### Morphological and biochemical properties of CRB14 isolate

With a high Cr (VI) tolerance and reduction ability the isolate CRB14 it was further subcultured in LB medium for morphological and biochemical characterization. In the LB agar plate, we observed off-white, circular phenotypic characteristics. After Gram staining and microscopic observation, it was found that the isolate is a gram-positive and rod-shaped bacteria. Biochemical reaction revealed that it is a motile and catalase-positive bacteria. In addition, it could utilize citrate and reduce nitrate. On the contrary, oxidase, urease, indole, methyl red, and Voges Proskauer test showed negative results. The bacteria also hydrolyzes starch and can ferment glucose and fructose. On the other contrary, it failed to ferment lactose and produce H2S gas.

### Formation of Biofilm

The isolate CRB14 was tested for biofilm formation cultured at 25, 50,100, and 200 mg/mL chromium VI. At the elevated level (200mg/ml of Cr(Vi)), the isolate produced weak biofilm (OD > ODc) according to the criteria of Stepanovic et al. (2007) [59].

### Analyses of plant growth-promoting abilities

We observed the plant growth-promoting properties in CRB14. This isolate fixed atmospheric N_2_ as it could grow in a nitrogen-free New Fabian Broth (Nfb) medium. The amount of nitrogen fixed by the isolate was 10.34 μg/ml. It was also able to solubilize insoluble phosphate in NBRIP media indicating its potential to promote the growth of plants under phosphate-limited conditions. In quantitative analysis, the isolate could solubilize phosphate 1.05μg/ml. The isolate in its respective culture of ∼10^8^ CFU/ml was screened for their ability to produce IAA. The IAA production was found to be 28.165 μg/ml. In continuation of addressing PGP activities, this study characterized the siderophore-producing ability of the test isolate. The siderophore-producing ability was scored qualitatively by the formation of orange zones produced around the bacterial growth. In a quantitative approach, it was observed that it can produce 49.02 psu (percent siderophore unit). Moreover, it degraded cellulose and showed the ability to produce ammonia (Supplementary Figure 02).

### Molecular Characterization

#### Plasmid and Cr-associated genes

The genetic determinants of resistance to heavy metals are often found in plasmids and transposons [60]. Therefore, we tested if there were any detectable plasmids present. Upon plasmid extraction and gel electrophoresis, isolate CRB14 showed no plasmid. This finding suggests that tolerance to Cr by CRB14 is possibly mediated by chromosomal genes.

To investigate whether CRB14 isolate possesses some known heavy metal tolerance genes, PCR amplification of specific chromium tolerant genes *chrA* (chromate transport protein), and *ycnD* (FMN reductase) was conducted. Analysis of the PCR product showed that the isolate produced a band of 268 bp size, which is the expected size for the *chrA* gene according to Patra et al. (2010) (Supplementary Figure 03) [61]. On the other hand, no band was visualized for the ycnD gene. This indicates that CRB14 contains chromium-resistant genes. The presence of *the* gene was further confirmed by sequencing the PCR products.

#### Genomic characterization of isolate CRB14

The aforementioned experimental research was further validated through comprehensive genome sequencing and analyses. The complete genome of CRB14 was assembled into 344 contigs, totaling 5,217,143 bp with an average GC content of 35.3%. A total of 5255 genes were predicted, including 2043 hypothetical proteins, 3 rRNA, 37 tRNA genes, and 1 tmRNA gene. Additional genome features are presented in the **Supplementary Table 01**. The circular view of the genome from PROKSEE online software is presented in **Figure 05** highlighting the GC distribution, and GC-skew of the genome. The genome data for CRB14 has been submitted to NCBI under Biosample accession number SAMN41810974 and BioProject ID PRJNA1123424.

**Figure 05:**
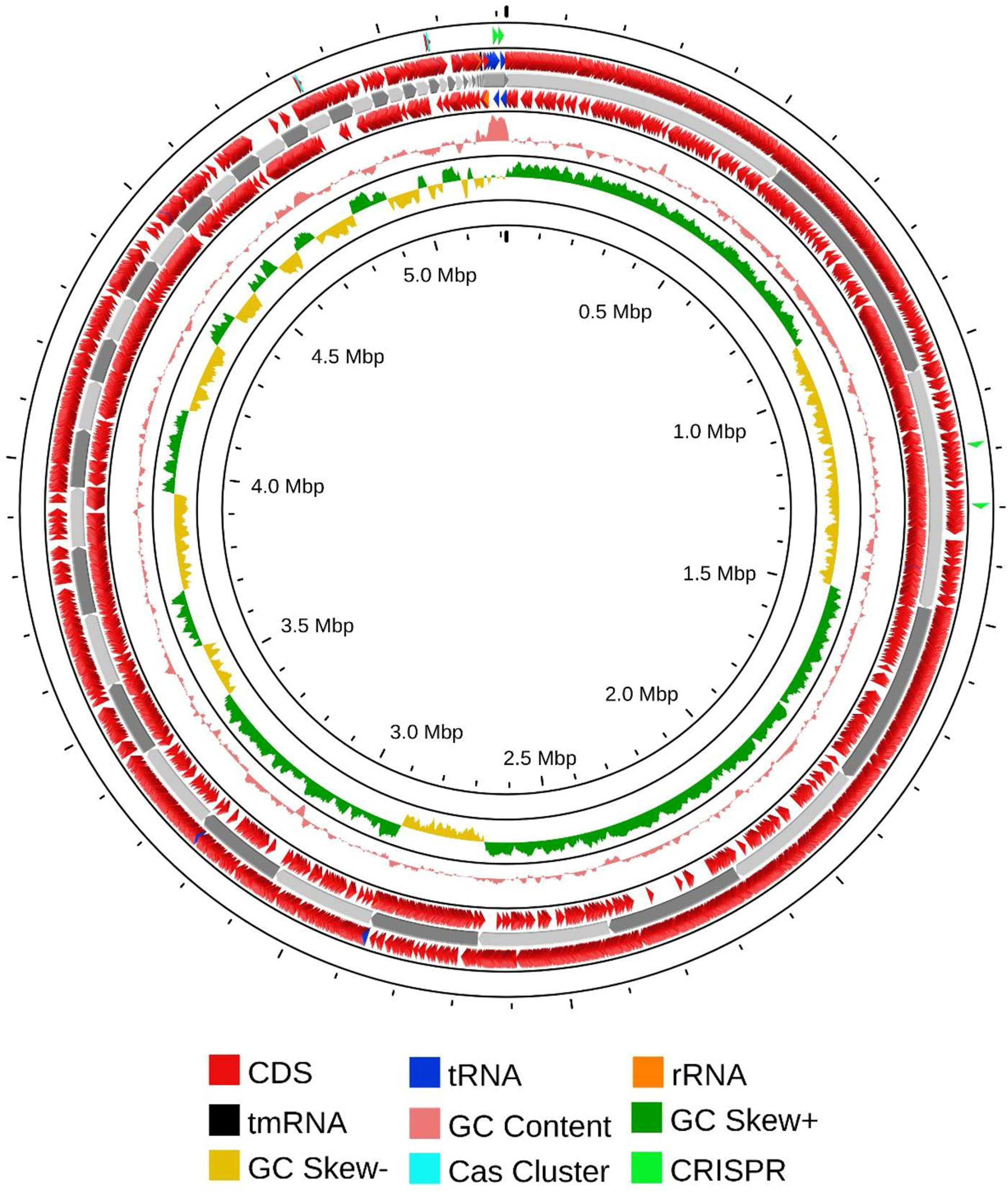
Nuclear genome circle diagram of CRB14. From outside to inside, coding genes (positive-sense strand), coding genes (negative-sense strand), tRNA (blue) and rRNA (orange), tmRNA (black), CRISPR (green), Cas cluster (cyan), GC ratio and GC-skew.

#### Genomic islands and CRISPR prediction

Genomic islands (GIs) are vital for microbial genome evolution as they harbor diverse genes that enhance adaptability to the surrounding environment. These regions influence key traits contributing to pathogenesis, antibiotic resistance, metabolism, etc. Thus, the prediction of GIs has become increasingly essential in microbial genome analysis [62]. 34 putative GIs were found in CRB14 and the size of GIs ranged from 5.7 to 38.9 kb. CRISPR guide RNAs can be tailored to target pathogen-specific virulence and chromosomal genes, allowing for the repurposing of CRISPR-Cas systems to combat bacteria rather than solely defending against external threats. Moreover, CRISPR-Cas9 technology has the potential to generate antibiotics that specifically target antimicrobial-resistant pathogens by precisely targeting their genetic sequences. Therefore, detecting CRISPR elements within microbial genomes is crucial for gaining insight into the dynamics of microbial communities [63]. 4 CRISPRs, 4 cas genes, and 2 cas clusters were involved in CRB14.

#### Heavy metals and antibiotic-resistance genes

Biochemical characterization of *Bacillus tropicus* CRB14 demonstrated its ability to grow in the presence of various heavy metals, including chromium, cadmium, cobalt, iron, zinc, lead, and arsenic. To further support these findings, the annotated genome of CRB14 was analyzed for the presence of heavy metal tolerance genes. The results revealed abundant genes associated with resistance to chromium, cadmium, cobalt, and zinc (**Table 02**). These genomic findings align closely with the biochemical data, reinforcing CRB14’s capacity to tolerate and survive in environments contaminated with heavy metals.

**Table 02:**
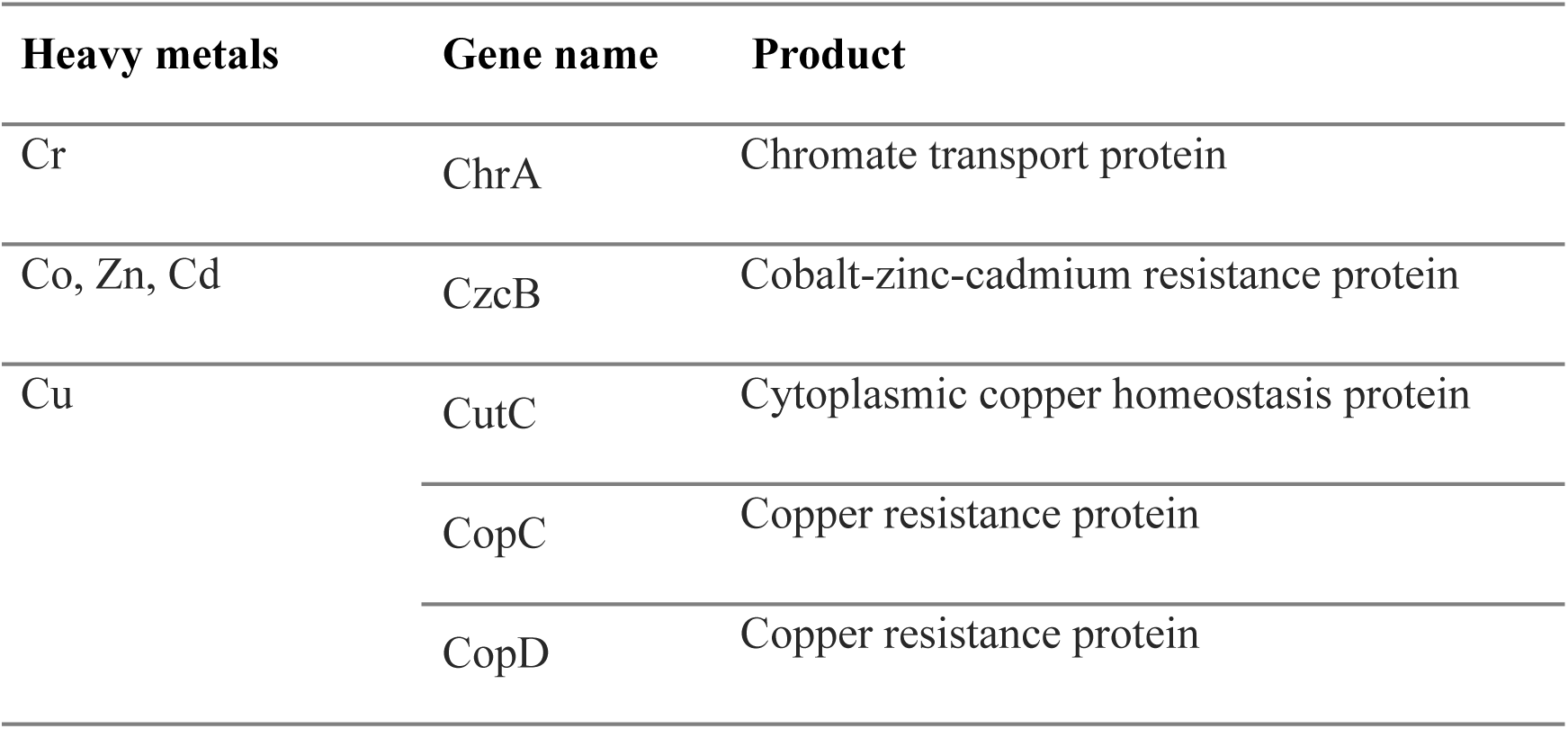
Genes related to heavy metal tolerance in the CRB14 genome.

Non-antibiotic compounds, such as antibacterial biocides and metals, frequently lead to the selection of bacterial strains that can also resist antibiotics. This dual resistance arises through two primary mechanisms: co-resistance, where resistance genes for both antibiotics and metals/biocides are located within the same cell, and cross-resistance, where a single mechanism (such as an efflux pump) provides resistance against both antibiotics and metals/biocides [64]. In light of this, the genome of CRB14 was evaluated for its resistance to antibiotics. The findings indicated that it was sensitive to several antibiotics, including vancomycin, teicoplanin, fosfomycin, tetracycline, and doxycycline, among others. In addition to the chromium resistance gene, it also harbors genes resistant to cobalt, zinc, cadmium, and copper. A detailed list of genes related to antibiotics and the associated resistance mechanisms is provided in **Supplementary Table 02**.

#### Genome comparison

In the BLAST search of the NCBI database using the 16S rDNA sequence of CRB14, numerous strains were identified. From these, 39 strains were selected for further analysis, as they exhibited 100% sequence similarity (E-value 0.00) to CRB14, meeting the species delineation threshold of 98.65% 16S rDNA sequence similarity [65]. The phylogenetic relationship of CRB14 depicted in **Figure 06**, shows its connection with closely related strains from GenBank. The genomic similarity between CRB14 and other strains was further assessed, enhancing the accuracy and resolution of the phylogenetic signals to demarcate bacterial species [66]. CRB14 exhibited an ANI value of 99.51% with *Bacillus tropicus* strain EMB20, leading to the identification of CRB14 as *Bacillus tropicus* (Supplementary Figure 04).

**Figure 06:**
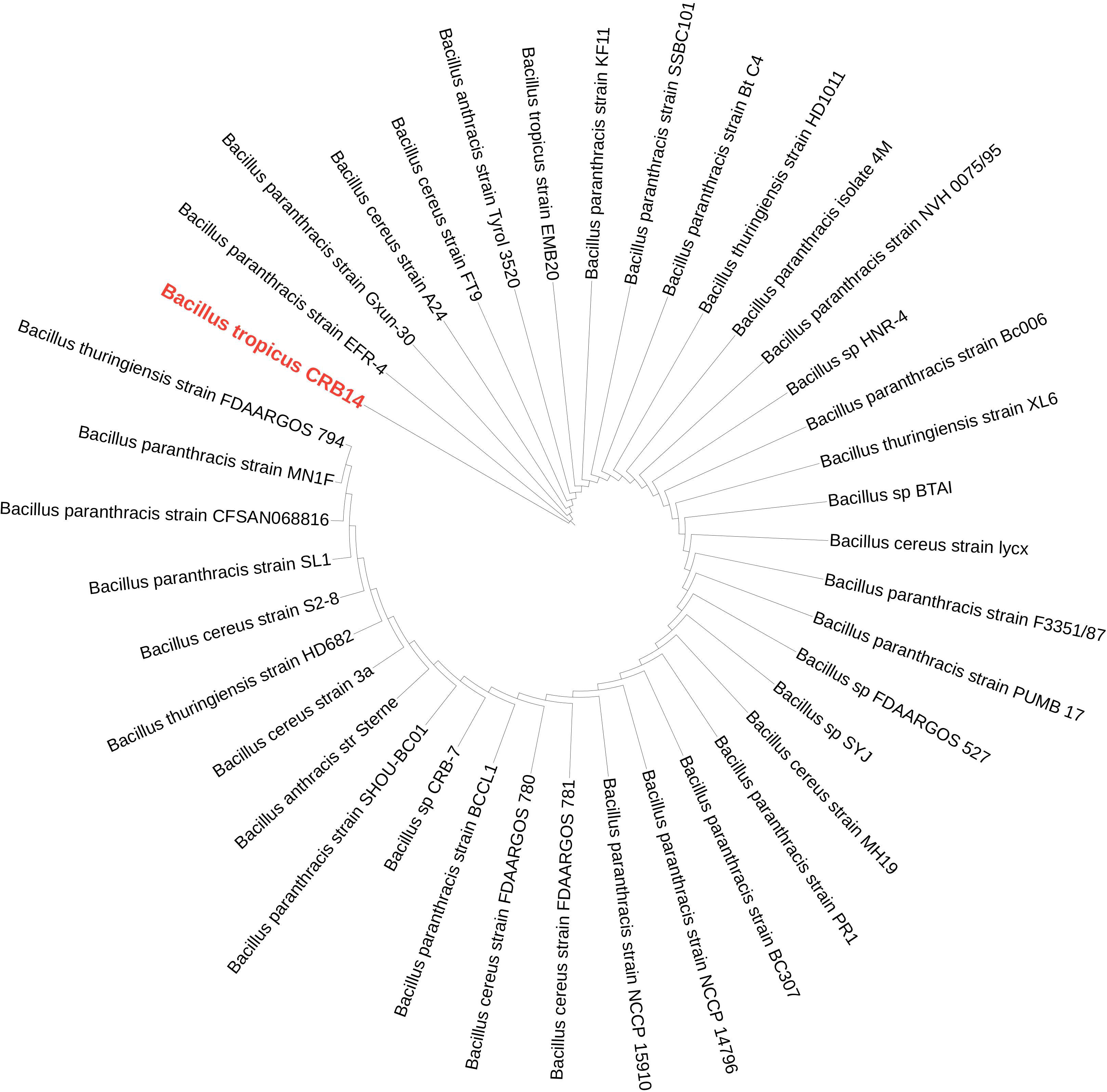
A maximum-likelihood phylogenetic tree based on 16S rRNA gene sequences of *Bacillus tropicus* CRB14 and other closely related strains. The isolate of interest is highlighted in red.

### Functional gene annotation

#### a) COG database annotation

The COG database categorizes proteins from fully sequenced genomes based on the principle of orthology [67]. To align all predicted CDS sequences of CRB14, the COG database was employed, utilizing the genome for functional and comparative analysis. Altogether 4,794 protein-coding sequences were successfully annotated into COG (**Figure 07**), categorized into 26 groups ranging from A to Z. A substantial number of genes were classified into the categories of “Amino acid transport and metabolism” (475), “Transcription” (452), “Inorganic ion transport and metabolism” (298), and “Carbohydrate transport and metabolism” (287). Meanwhile, genes associated with “Signal transduction mechanisms” (222), “DNA replication, recombination, and repair” (174), and “Secondary metabolites biosynthesis, transport, and catabolism” (102) were also present in notable proportions. The number of genes in the S (Function unknown) group is enormous. However, the current database failed to clarify their biological functions, indicating that many unknown functions are worthy of further elucidation.

**Figure 07:**
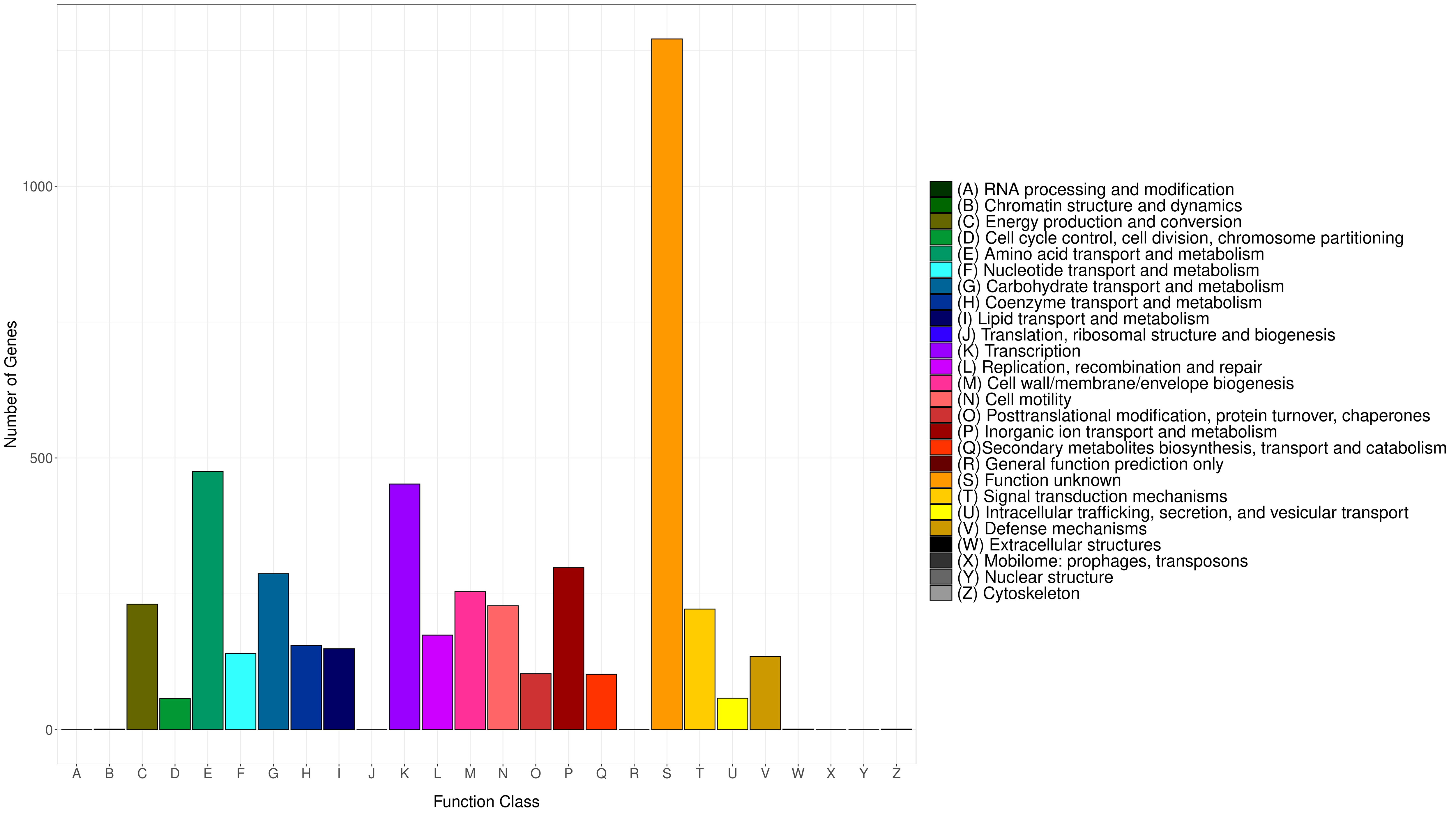
Bar graph demonstrating the COG classifications of the genome CRB14. Each bar corresponds to a specific classification, highlighting these proteins’ diverse roles in bacterial metabolism and physiological processes. The abscissa represents the various COG categories, while the ordinate shows the number of genes assigned to each category.

#### b) KEGG database annotation

The KEGG database connects molecular functions of genes and proteins to ortholog groups, aiming to relate gene sets in the genome to the higher-level functions of cells and organisms [68]. In total, 2,686 genes were classified into 41 pathways in the KEGG database and arranged into eight fundamental classes, each comprising several subclasses (**Figure 08**). Most genes fell into the following categories: under “metabolism”, there were 289 genes involved in carbohydrate metabolism, 258 in amino acid metabolism, 162 in the metabolism of cofactors and vitamins, and 133 in energy metabolism; under “environmental information processing”, 121 genes were related to membrane transport and 107 to signal transduction; and under “genetic information processing”, 81 genes were associated with translation and 70 with replication and repair.

**Figure 08:**
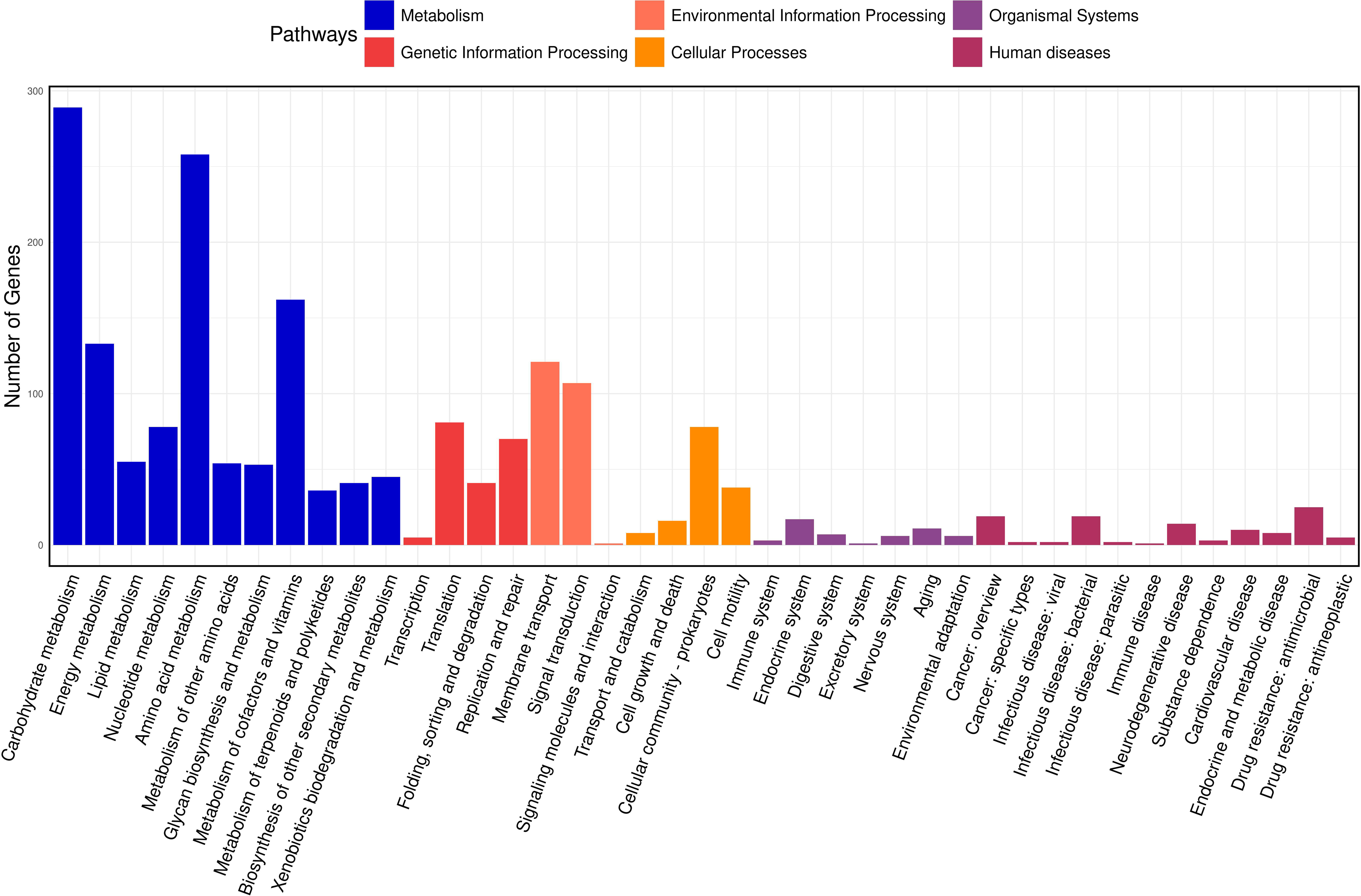
KEGG classification of the predicted coding sequences. The x-axis denotes the various pathways, and the y-axis indicates the number of genes assigned to each pathway. The bars are color-coded according to the six major pathway classes, indicating that the majority of genes are involved in metabolism.

#### c) GO Database Annotation

Gene Ontology (GO) is a structured vocabulary project designed to describe the functions of gene products across all organisms. It categorizes molecular and cellular biology into three domains: molecular function, cellular component, and biological process, each with its subdomains [69]. According to the GO functional annotation results for CRB14, the cellular components domain was predominantly represented by genes related to intracellular anatomical structures and the cytoplasm. Similarly, the biological process category revealed that most genes were associated with cellular metabolic processes and organic substance metabolism. In the molecular function domain, the most prevalent functions were binding, ribosomal structural components, and catalytic activities. **Figure 09** depicts the most significant subdomains within each major domain of the GO database.

**Figure 09:**
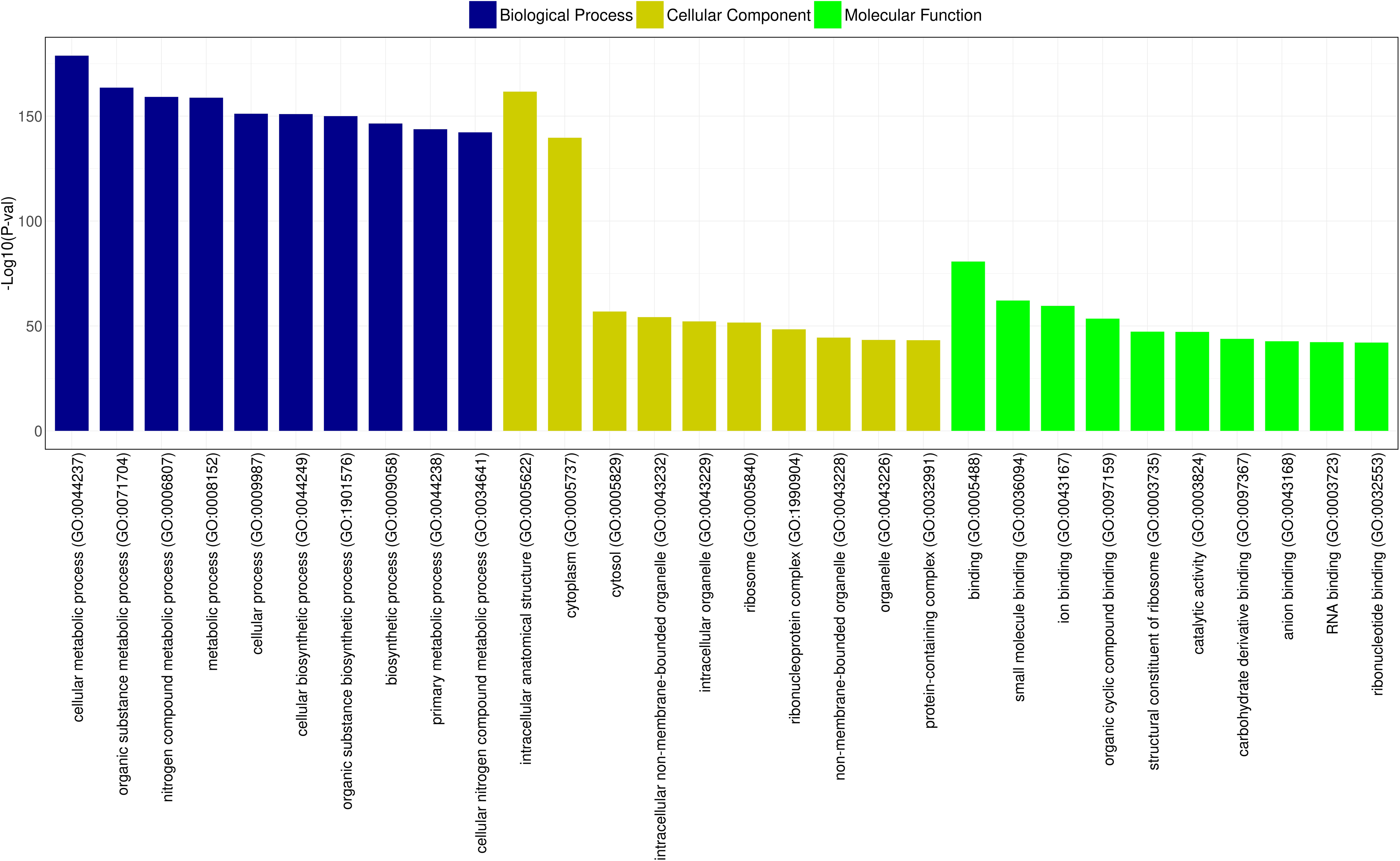
GO functional classification of CRB14. The x-axis shows the GO categories, and the y-axis represents the -log10 (P-value) for the top 10 terms in biological process (blue), cellular component (yellow), and molecular function (green).

### Predictive analysis of biosynthetic gene clusters (BGCs)

AntiSMASH predicted a total of nine gene clusters for secondary metabolite synthesis: Tarpene, RRE-containing, Betalactone, RiPP-like, Linear azol(in)e-containing peptides(LAP), NRPS-independent, IucA/IucC-like siderophores (NI-siderophore), Non-ribosomal peptide metallophores (NRP-metallophore), Class II lanthipeptides like mutacin II (U40620). There are three strictness levels for cluster detection: strict, relaxed, and loose. RRE-containing and RiPP-like clusters are in a relaxed level and the rest of them in a strict level. All these groups of genes included genes involved in additional biosynthetic, core biosynthetic, transport-related, regulatory, and other genes (**Figure 10**), in which siderophore and NRP-metallophore have been extensively studied. These are the most effective ways for microorganisms to take up iron from iron-poor environments: bacteria containing siderophores use specific ATP-dependent membrane-associated transporters to deliver the Fe(III)-siderophore complex to the cell. Inside the cell, Fe(III) is enzymatically reduced to soluble and solid-phase Fe(II), which quickly transfers its electron to Cr (VI), becoming the reductant for Cr (VI) [22]. The biosynthetic gene cluster family (BiG-FAM) database is used to be shown with the summary of all best BGC-to-GCF pairings with a distance lower than 900 (original threshold value) depicting a good match to at least one GCF that belongs to the genus *Bacillus* in the database and the completeness of the gene (**Supplementary Table 03**).

**Figure 10:**
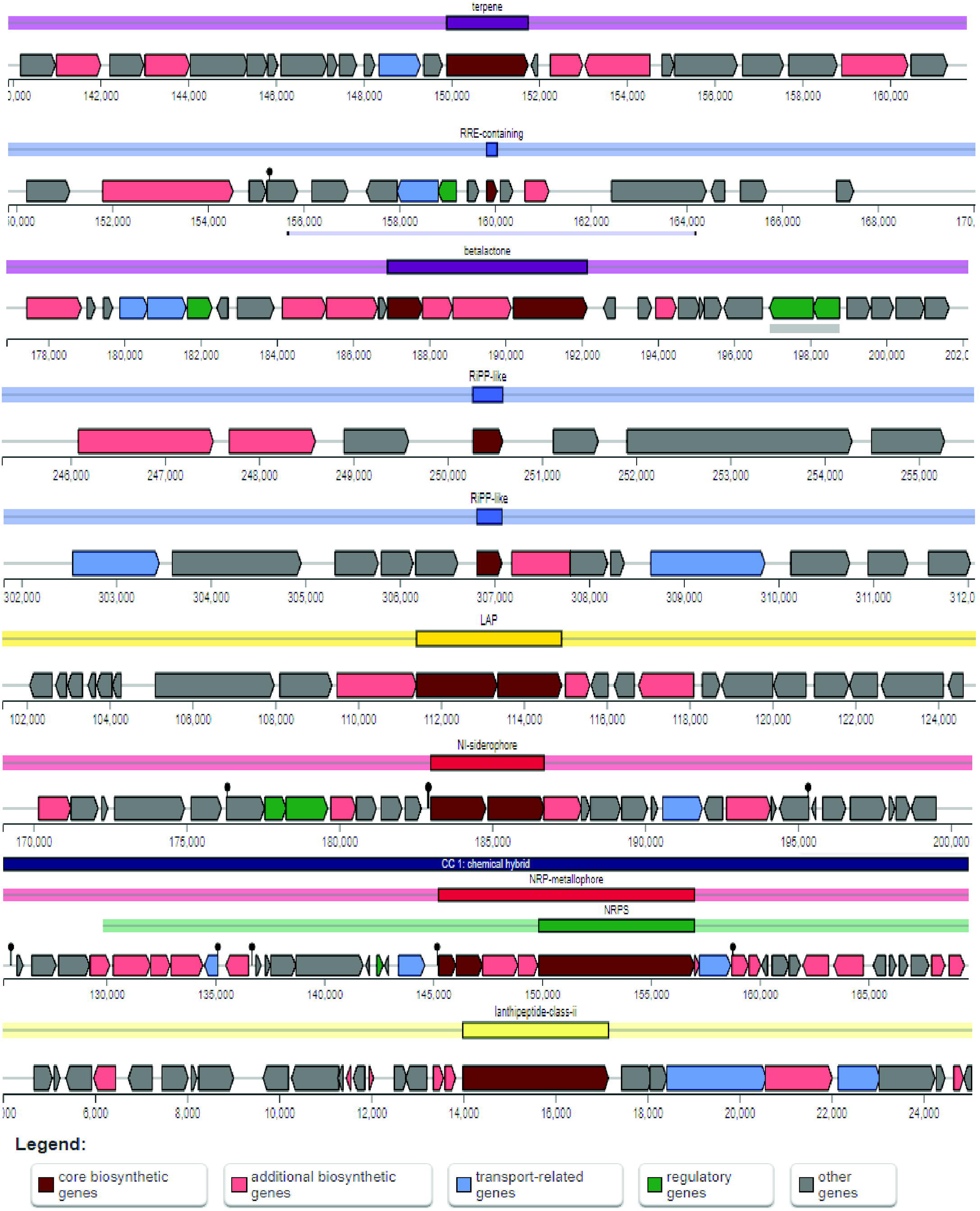
Schematic diagram of nine secondary metabolite biosynthetic gene clusters in *B. tropicus* CRB14. Potential secondary metabolite biosynthetic gene clusters were predicted using antiSMASH. Color-coded blocks indicate different gene functions: dark red for core biosynthetic genes, light red for additional biosynthetic genes, blue for transport-related genes, green for regulatory genes, and gray for other genes.

## Discussion

Widening the understanding of heavy metal detoxifying bacteria is crucial for their utilization for bioremediation purposes. Moreover, evidence-based information potentially acts as a source of genetic engineering. This study investigated a promising bacterial isolate, CRB14, showcasing its excellent tolerance and reduction capabilities towards chromium (VI), a toxic heavy metal contaminant.

The isolate demonstrated an exceptional tolerance to chromium (VI), thriving in concentrations as high as 5000 mg/L in agar and 900 mg/L in broth. This tolerance is particularly noteworthy as chromium (VI) is highly toxic to most microorganisms. Furthermore, CRB14 exhibited significant chromium (VI) reduction ability, achieving over 86% reduction within 96 hours at higher concentrations. This reduction capability is crucial for bioremediation strategies as it transforms chromium (VI) into its less toxic form, chromium (III). Previously reported, *Bacillus cereus* isolate PGBw4 mitigated toxic effects of Cr(VI) more efficiently from 100mg/L to 500mg/L in LB broth and *Bacillus amyloliquefaciens* has tolerance to Cr(VI) up to 860 mg/L in LB broth. [70] [71] In addition, it is found that presumptively identified *Bacillus sp.* can tolerate 7700 mg/L Cr (VI) in LB agar media and can reduce 1000mg/l Cr (VI) in LB broth media. However, their reduction efficiency is only 44.33% in 500mg/l Cr (VI) in LB broth media [72]. It is noteworthy that Cr reduction did not completely relate to its tolerance level. Hence, it would be wise to select isolates for remediation purposes based on their Cr(VI) reduction capability rather than their high tolerance to Cr(VI) [73].

Besides chromium, CRB14 demonstrated resistance to an array of heavy metals, including cadmium, cobalt, iron, zinc, lead, and arsenic, highlighting its potential for broader application in environments contaminated with multiple metals. Notably, the isolate failed to grow in the presence of nickel and mercury, suggesting the presence of selective metal resistance mechanisms. Nature can address heavy metal pollution through microorganisms that adapt to contaminated environments. These microbes develop certain resistance mechanisms and often form biofilms, which help in metal sequestration and remediation by secreting exopolysaccharides (EPS) that bind positively charged metal ions and produce enzymes and biosurfactants [74], [75]. While the exact effects of heavy metals on biofilms are not fully understood, some studies have demonstrated the ability of biofilms to remove Cr(VI). For example, biofilms formed by Pseudomonas putida, Arthrobacter viscosus, or mixed microbes from activated sludge have shown Cr(VI) reduction and immobilization [76]. In our study, although biofilm formation was limited under elevated chromium levels, its presence could enhance the survival of CRB14 in harsh environments with high metal concentrations [59].

According to recent research, microorganisms suitable for bioremediation may promote plant growth. This is attributed to their abilities to detoxify and degrade toxins, as well as their production of growth-promoting metabolites. Our study revealed several plant growth-promoting (PGP) properties of CRB14. These include nitrogen fixation (10.34 μg/ml) and positive for ammonia production, cellulase production, phosphate solubilization (1.05 μg/ml), IAA production (28.165 μg/ml), and siderophore production(49.02 psu). IAA is a plant hormone that stimulates root growth and development. Nitrogen fixation and ammonia production by CRB14 can enrich the soil with nitrogen, a vital nutrient for plant growth. Phosphate solubilization by CRB14 can increase the availability of phosphorus in the soil, another essential plant nutrient. Furthermore, CRB14 demonstrated siderophore production, which can help plants acquire iron, a vital nutrient for growth [77]. These iron acquisition systems might indirectly contribute to Cr(VI) reduction. On the other hand, CRB14 has cellulose degradation ability. Plants typically contain up to 60% cellulose, the decomposition of cellulose is a key activity of soil bacteria and is vital to the energy flow through soils and the cycling of N, P, and S, where immobilization generally accompanies cellulose decomposition [78]. These PGP traits suggest CRB14’s potential as a biofertilizer that can enhance plant growth and metal stress tolerance.

The genetic determinants of resistance to heavy metals are often found in plasmids and transposons [60]. Plasmid analysis revealed no detectable plasmids in CRB14, suggesting chromosomal residence of Cr resistance genes. A similar result is also shown in *Bacillus spp* [79]. By analyzing PCR amplification products, it was revealed that the Cr-tolerance-related gene ChrA exists in the genome of CRB14. Chr-A encodes transporters involved in the transport of Cr(III) and Cr(VI) through functions such as creating a transmembrane proton gradient, generating membrane potential, and facilitating electron transfer [80]. Upon exposure to chromium stress, Cr(VI) enters bacterial cells via sulfate ion channels and is carried out by ChrA transmembrane proteins, henceforth, mitigating the harmful effects of hexavalent chromium [22], [61]. Conversely, the absence of the ycnD gene, associated with FMN reductase, suggests alternative pathways for Cr(VI) reduction might be present in the genome of CRB14.

In support of the observational findings, the genome of CRB14 was thoroughly analyzed, revealing the presence of chromium and several other heavy metal resistance genes. This provides compelling evidence for the chromosomal location of these resistance genes in the isolate. The whole-genome analysis also uncovered 34 putative genomic islands (GIs) within it’s genome. GIs play a crucial role in the dissemination of heavy metal resistance genes among bacteria, enhancing their adaptability in metal-contaminated environments [81]. Researchers have shown that these discrete DNA segments, acquired via horizontal gene transfer (HGT), significantly contribute to bacterial genome plasticity by integrating beneficial genetic material in response to environmental pressures [82]. Through HGT, GIs may harbor genes involved in detoxifying or resisting toxic metals, offering a survival advantage in metal-rich environments. In turn, this genetic exchange promotes environmental adaptation and can result in the evolution of highly resistant bacterial strains capable of thriving under extreme conditions.

Functional annotation implies the involvement of diverse metabolic pathways and the potential utilization of siderophores for Cr resistance. This knowledge could contribute to the development of bioremediation strategies for chromium-contaminated environments. Future research is to be focused on the functional characterization of the identified Cr resistance genes paving the way for its potential application in genetic engineering for bioremediation.

## Conclusion

This study presents the characterization of a chromium-reducing bacterium, *Bacillus tropicus* CRB14, collected from tannery effluent. Both experimental data and genome analysis revealed the presence of several genetic determinants related to resistance against chromium and other heavy metals, including cadmium, cobalt, iron, lead, arsenic, and zinc. Furthermore, CRB14 exhibited strong plant growth-promoting capabilities, such as efficiently solubilizing inorganic phosphate, producing siderophores to enhance nutrient uptake, and synthesizing indole-3-acetic acid (IAA) to stimulate plant growth. To the best of our knowledge, this is the first report offering new insights into chromium and multi-metal resistance in this species of *Bacillus*. Future research will further elucidate the mechanisms and pathways involved in CRB14’s multi-metal resistance, potentially supporting its application in heavy metal-polluted environmental and agricultural sites.

## Supplementary Files

**Supplementary Figure 01:** Optimization of physical factors (pH, temperature, shaking speed) for the growth of CRB14. After incubation, the bacteria exhibited the best growth at a) 35°C with b) 150 rpm shaking at c) pH 8.

**Supplementary Figure 02:** Plant growth promoting (PGP) activities determination of the isolate CRB14.

**Supplementary Figure 03:** Molecular Characterization of CRB14. a) Plasmid profiling of CRB14 shows no bands in the gel, indicating the absence of plasmids in the isolate. b) PCR amplification of Cr(VI) Resistant chrA and ycnD Gene. Lane 1 and Lane 2 represent the PCR products using chrA and ycnD gene-specific primers, respectively. A band is visible in Lane 1, confirming the presence of the chrA gene, while no band is observed in Lane 2, indicating the absence of the ycnD gene in the CRB14 genome.

**Supplementary Figure 04:** Species identification based on whole-genome sequencing. A heat map illustrating the similarity levels based on the average nucleotide identities (ANI) of the whole genomes of 40 related strains. ANI percentages are visualized with color intensity — red for higher similarity and blue for lower. The two most similar species, *Bacillus tropicus* strain EMB20 and *Bacillus tropicus* CRB14, are highlighted within a box.

---

**Supplementary Table 01:**
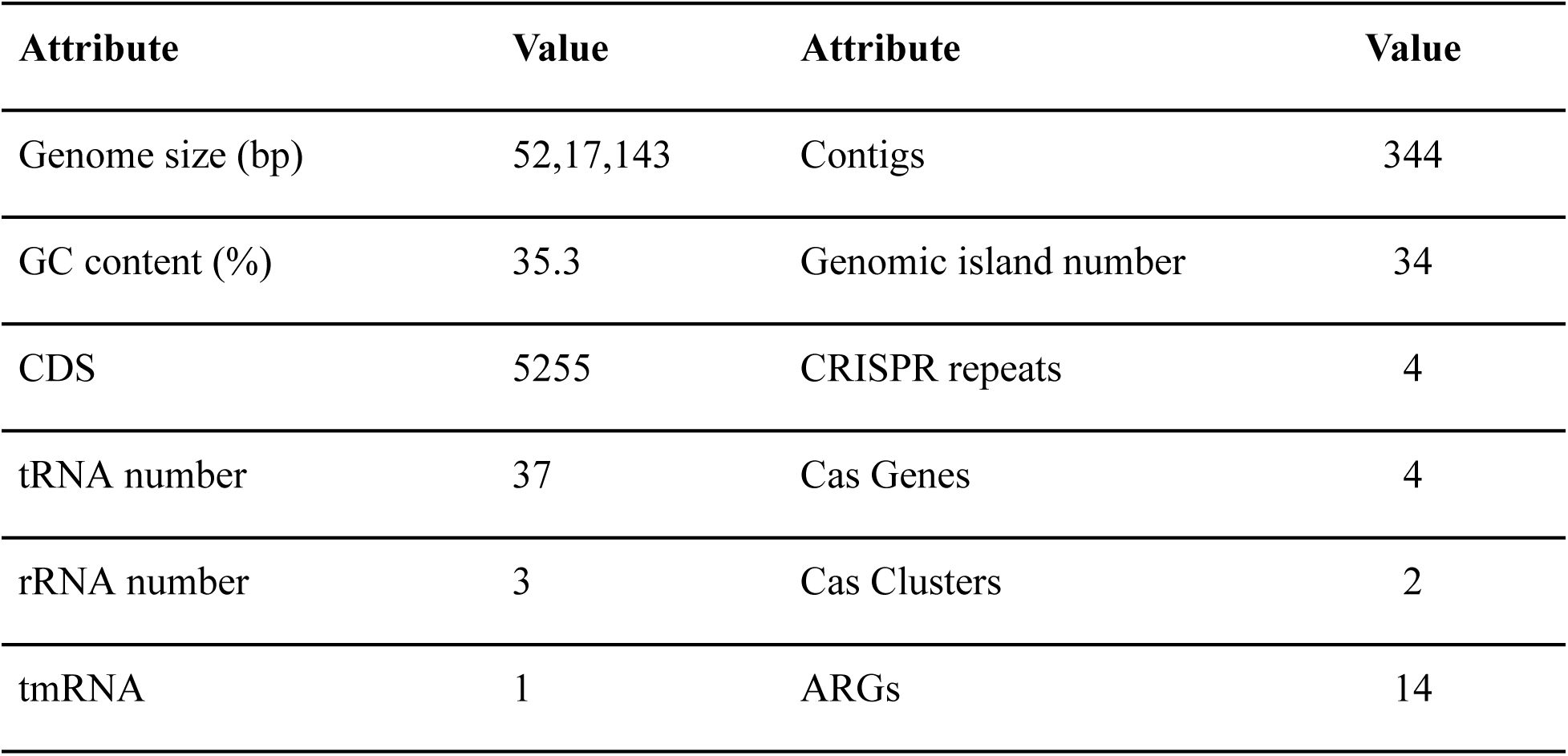
General features of the isolate CRB14.

**Supplementary Table 02:**
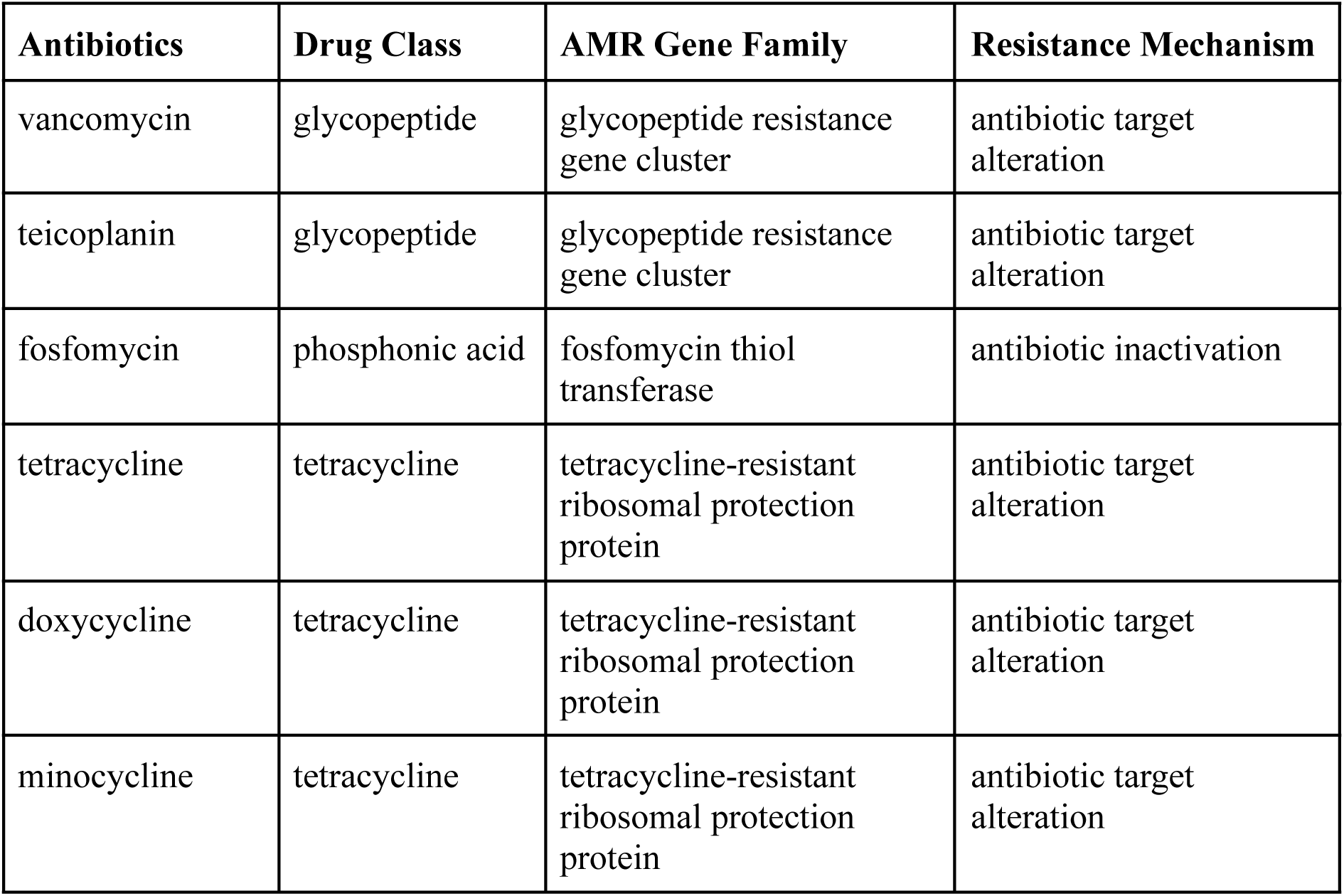
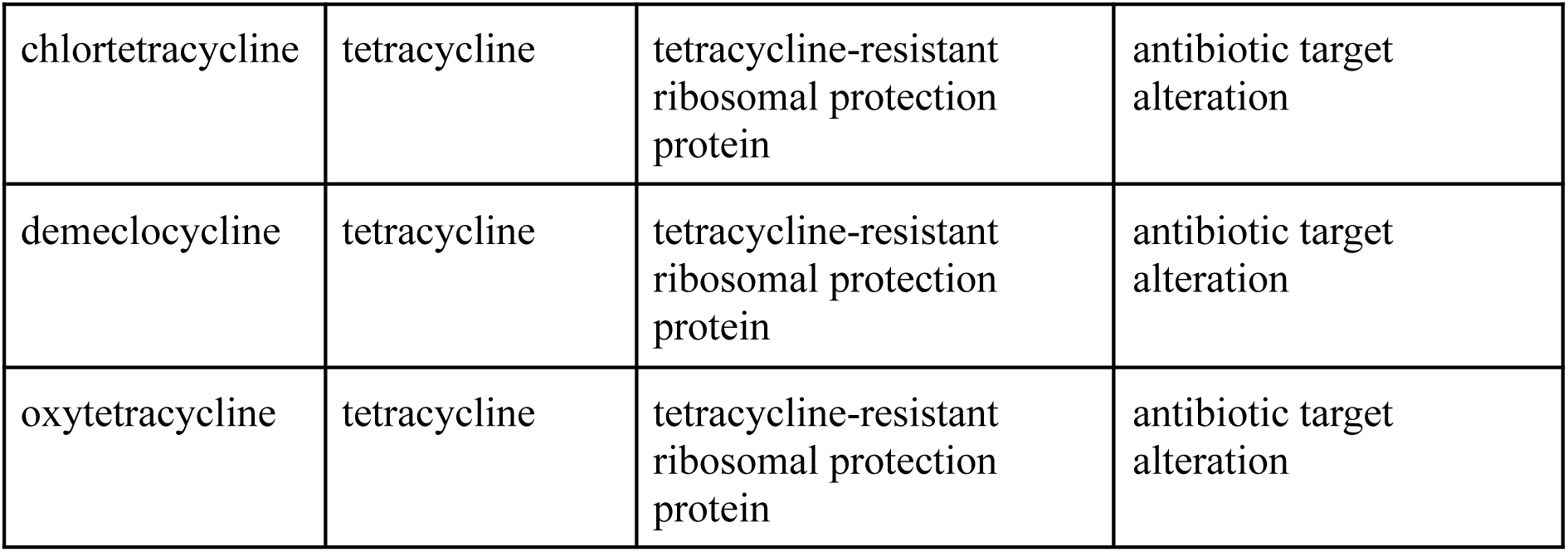
List of antibiotics and associated resistance mechanism found in CRB14.

**Supplementary Table 03:**
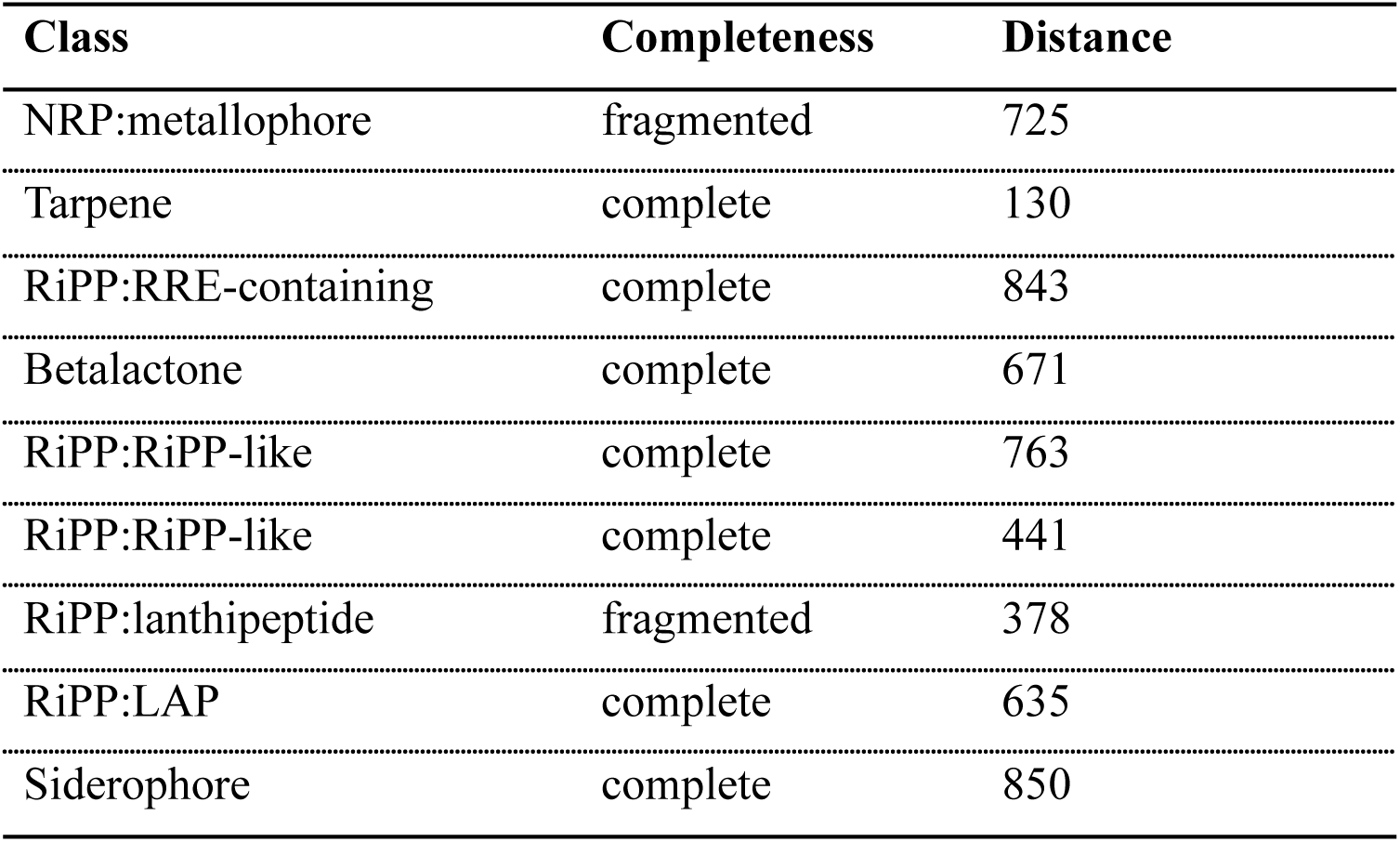
Biosynthetic gene cluster family summary of all best BGC-to-GCF pairings.

