## Supplementary figures and images for "Characterization and molecular insights of a chromium-reducing bacterium *Bacillus tropicus*"

### Supplementary Figure 1

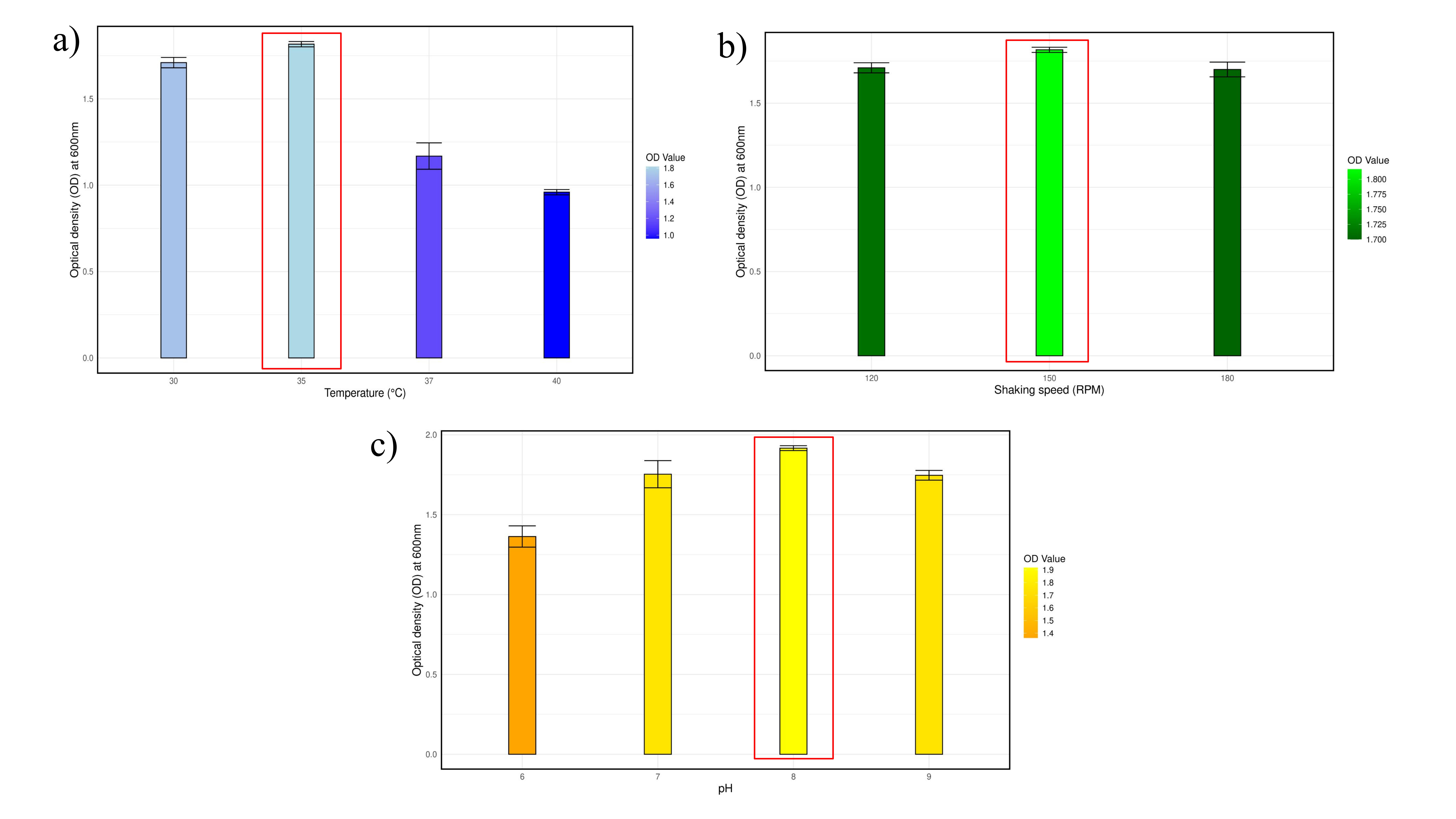

### Supplementary Figure 2

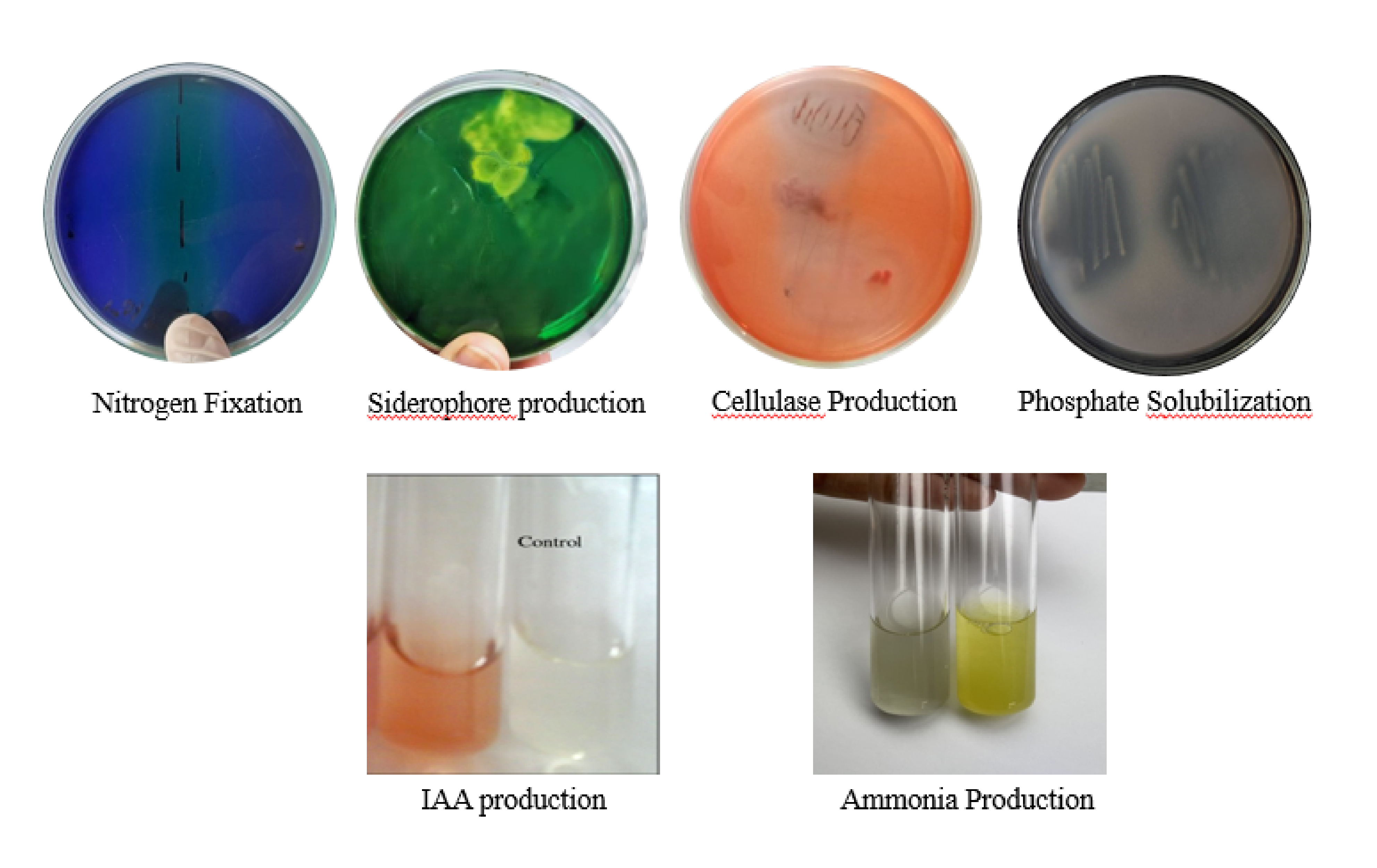

### Supplementary Figure 3

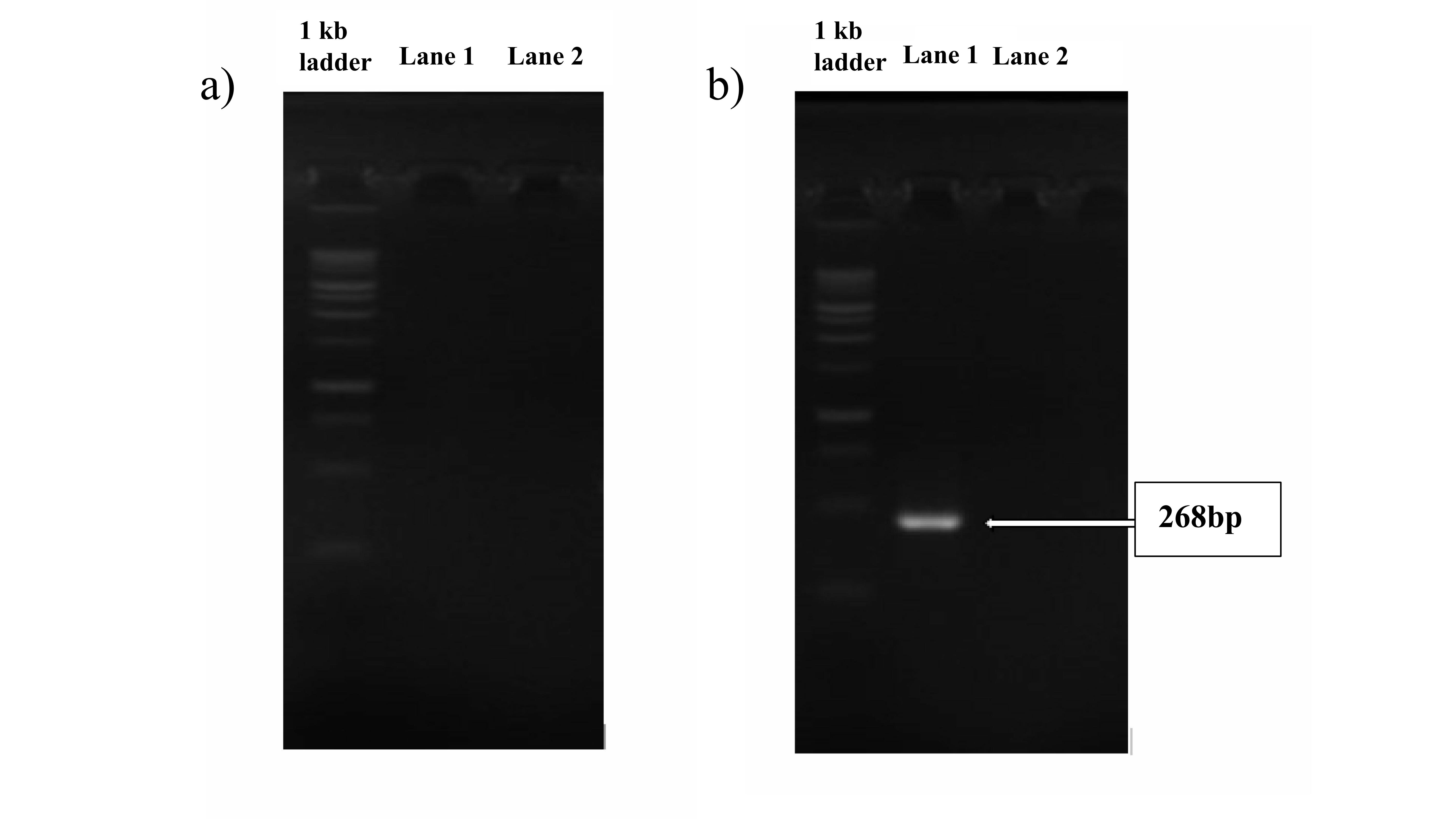

### Supplementary Figure 4

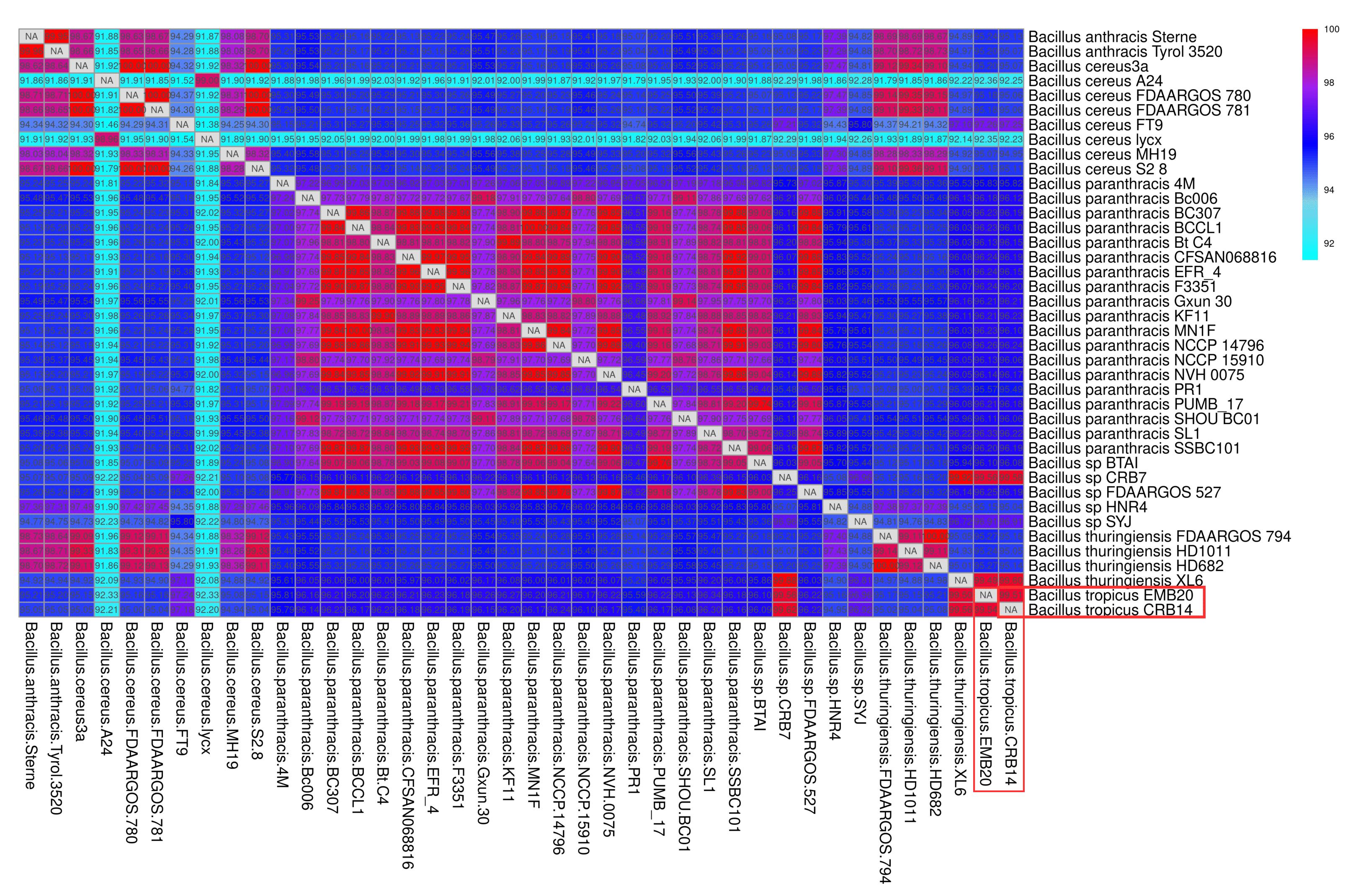
